# The interaction of aging and oxidative stress contributes to pathogenesis in mouse and human Huntington disease neurons

**DOI:** 10.1101/800268

**Authors:** Emily Machiela, Ritika Jeloka, Nicholas S. Caron, Shagun Mehta, Mandi E. Schmidt, Colton M. Tom, Nalini Polturi, Yuanyun Xie, Virginia B Mattis, Michael R. Hayden, Amber L. Southwell

## Abstract

Huntington disease (HD) is a fatal, inherited neurodegenerative disorder caused by a mutation in huntingtin (*HTT*). While mutant HTT is present ubiquitously throughout life, HD onset typically occurs in mid-life. Oxidative damage accumulates in the aging brain and is a feature of HD. We sought to interrogate the roles and interaction of age and oxidative stress in HD using primary Hu97/18 mouse neurons, neurons differentiated from HD patient induced pluripotent stem cells (iPSCs), and mice. We find that primary neurons must be matured in culture for canonical stress responses to occur. Furthermore, when aging is accelerated in mature HD neurons, mutant HTT accumulates and sensitivity to oxidative stress is selectively enhanced. Furthermore, we observe HD-specific phenotypes in iPSC-derived neurons and mouse brains that have undergone accelerated aging. These findings suggest a role for aging in HD pathogenesis and interaction between biological age of HD neurons and sensitivity to exogenous stress.

## Introduction

Huntington disease (HD) is an autosomal dominant neurodegenerative disease caused by an expanded polyglutamine encoding CAG tract in the *Huntingtin* (*HTT*) gene (MacDonald, 1993). HD neuropathology originates with degeneration of striatal medium spiny neurons (MSNs), followed by white matter loss and widespread forebrain atrophy. HD is characterized by progressive movement, psychiatric, cognitive, and behavioral abnormalities, with motor symptoms typically beginning between ages 35-45 (Bates et al., 2015), and death occurring 10-20 years after symptom onset. Currently, there are no disease modifying therapies for HD.

Age-of-onset of HD is negatively correlated with CAG tract length (Andrew et al., 1993; Snell et al., 1993) However, this only explains ∼50-70% of variation in disease onset age, with other genetic and environmental factors making up the remainder (Wexler et al., 2004). Patients with the same expanded CAG tract length can have disease onset up to 50 years apart. Additionally, those with identical mutations can progress at different rates. This suggests that factors aside from the CAG repeat can have a strong influence on disease pathogenesis.

A genome-wide association study (GWAS) conducted in HD patients has identified single nucleotide polymorphisms (SNPs) in genes associated with early and late disease onset (Genetic Modifiers of Huntington’s Disease, 2015). Several loci that impact disease onset were identified. A pathway analysis of these loci revealed the strongest signals were found in genes involved in DNA damage repair. Specifically, these genes are responsible for pathways that repair breaks in DNA associated with damage from stressors, such as oxidative stress. More recently, a GWAS was performed to identify factors that influence progression of HD. Similar to findings for age of onset, the SNPs with the strongest signals identified to influence progression of HD were found in genes recruited to DNA after oxidative DNA damage, for base-excision repair (Lai et al., 2016; Moss et al., 2017).

In HD, there is evidence for both reduced oxidative stress responses and increased oxidative damage in neurons (Johri and Beal, 2012; Kovtun et al., 2007; Rotblat et al., 2014; Sepers and Raymond, 2014). Reactive oxygen species (ROS) are byproducts of normal cellular metabolism, cleared primarily by endogenous antioxidant systems in the cell (Beckhauser et al., 2016). Oxidative stress occurs when oxidants overwhelm antioxidants, which leads to oxidative damage to proteins, nucleic acids, and other biomolecules. Increased markers of oxidative damage such as protein carbonylation, lipid peroxidation, and DNA damage have been found in postmortem HD brains and the brains of HD mice (Bogdanov, Andreassen, Dedeoglu, Ferrante, & Beal, 2001; Browne & Beal, 2006; Browne, Ferrante, & Beal, 1999). Additionally, in the blood of HD patients, decreased antioxidant protein concentration and increased oxidative damage have been found (Chen et al., 2007; Klepac et al., 2007), suggesting that the impaired oxidative stress response occurs in both the brain and peripheral tissues of HD patients. The consequences of oxidative damage include loss of protein or organelle function, and permanent damage or failed repair of DNA and RNA. Oxidative damage can also increase somatic expansion of the CAG repeat tract in HD neurons, a phenomenon associated with earlier HD onset (Jonson et al., 2013; Kovtun et al., 2007)

While a small percentage of HD patients experience the juvenile onset form of the disease (Nance and Myers, 2001), HD is primarily a disease of aging; patients live with the mutation for many decades before symptom onset occurs. One difficultly in modeling a late-onset disease such as HD is the lack of age in typical model systems. Rodents have relatively short life-spans, and most studies are performed in young animals for time and cost efficiency. Moreover, both rodent primary neurons and neurons derived from patient induced pluripotent stem cells (iPSCs) are embryonic in nature, possessing no markers of aging. Embryonic neurons from adult-onset HD patients are robust with few phenotypes (Consortium, 2012; V. B. Mattis et al., 2015), thus not an appropriate model of HD. Additionally, oxidative and DNA damage as well as somatic expansion, potential drivers of HD pathogenesis, increase with aging. Combined with evidence of advanced biological age in HD brains (Grima et al., 2017; Horvath et al., 2016), age-related defects in the nuclear pore complex of HD neurons (Grima et al., 2017), and the fact that aging can render MSNs to mutant HTT (mtHTT) toxicity (Diguet et al., 2009) this suggests an interplay between age, oxidative stress, and mutant HTT in HD neurodegeneration and thus onset and/or progression.

It has been shown that progerin, a truncated form of lamin A that accumulates during natural aging (McClintock et al., 2007) and causes the accelerated aging disease Hutchinson-Gilford Progeria Syndrome when overproduced by a mutation in lamin A (Eriksson et al., 2003), can be used to age iPSC-derived neurons. Progerin-induced aging was found to uncover disease-specific phenotypes in iPSC-derived neurons from Parkinson’s disease patients (Miller et al., 2013). This provides a way to study relevant cell types at relevant biological age for late onset diseases such as HD.

In this work, we sought to determine if age and oxidative stress can drive HD pathogenesis in neurons. We investigated both basal levels of oxidative damage and the response to exogenous oxidative stress in immature and mature primary cortical neurons from HD mice. Additionally, we investigated if inducing aging *in vitro* in HD primary cortical neurons as well as HD patient iPSC-derived neurons and *in vivo* in HD mice could uncover more robust HD-like phenotypes in these models.

These studies serve to provide new insight into HD modeling and disease mechanisms as well as to identify and validate therapeutic targets and strategies.

## Results

### Exogenous oxidative stress does not induce ROS in immature HD neurons

To induce oxidative stress in primary cortical neurons from Hu97/18 HD and Hu18/18 control mice, we treated neurons with hydrogen peroxide (H_2_O_2_), which is taken up by neurons and increases reactive oxygen species (ROS) (Whittemore, Loo, Watt, & Cotman, 1995) or neurobasal complete (NBC) vehicle. To ensure H_2_O_2_ was inducing oxidative stress, we first measured ROS 24h post-treatment. We found that H_2_O_2_ induced ROS in control neurons (**Figure 1A;** 2 way analysis of variance (ANOVA) treatment p=0.0021, genotype 0.8977, Bonferroni post-test different from NBC Hu18/18: 50µM p=0.0773; 100µM p=0.0133; 150µM p=0.0227), but failed to induce ROS in HD neurons (Different from NBC Hu97/18 18 50µM p=0.6155; 100µM p=0.9986; 150µM p=0.0390). To determine whether this unexpected phenomena was H_2_O_2_ specific, we treated neurons with menadione and staurosporine, two compounds that have also been shown to increase ROS in neurons (Kruman, Guo, & Mattson, 1998; Loor et al., 2010). We found a similar affect in menadione-treated neurons, with induction of ROS in control, but not HD, neurons (**Figure 1B**, treatment p=0.0196, genotype p=0.4051, post-test vs. PBS same genotype Hu18/18: p=0.0050; Hu97/18: p>0.9999). However, menadione treatment also caused nuclear abnormalities in neurons in the absence of elevated cell death (**Figure S1A**,**B**). Treatment with staurosporine, a non-selective protein kinase inhibitor, failed to induce ROS in control or HD neurons (**Figure 1C**, treatment p=0.2258, genotype p=0.0090, post-test vs.DMSO same genotype Hu18/18: p=0.3554; Hu97/18: p>0.9999). It is interesting to note that although staurosporine did not induce ROS in neurons, it did cause cell death selectively in HD neurons at 1µM concentration (**Figure S1C**, post-test different from NBC Hu97/18, 1000 nM p<0.0001, all other comparisons ns), suggesting that high levels of kinase inhibition are not well-tolerated by HD neurons. Due to the nuclear abnormalities observed with menadione treatment and the lack of ROS-induction with staurosporine treatment, we have used H_2_O_2_ treatment to induce oxidative stress in all subsequent experiments.

**Figure 1.**
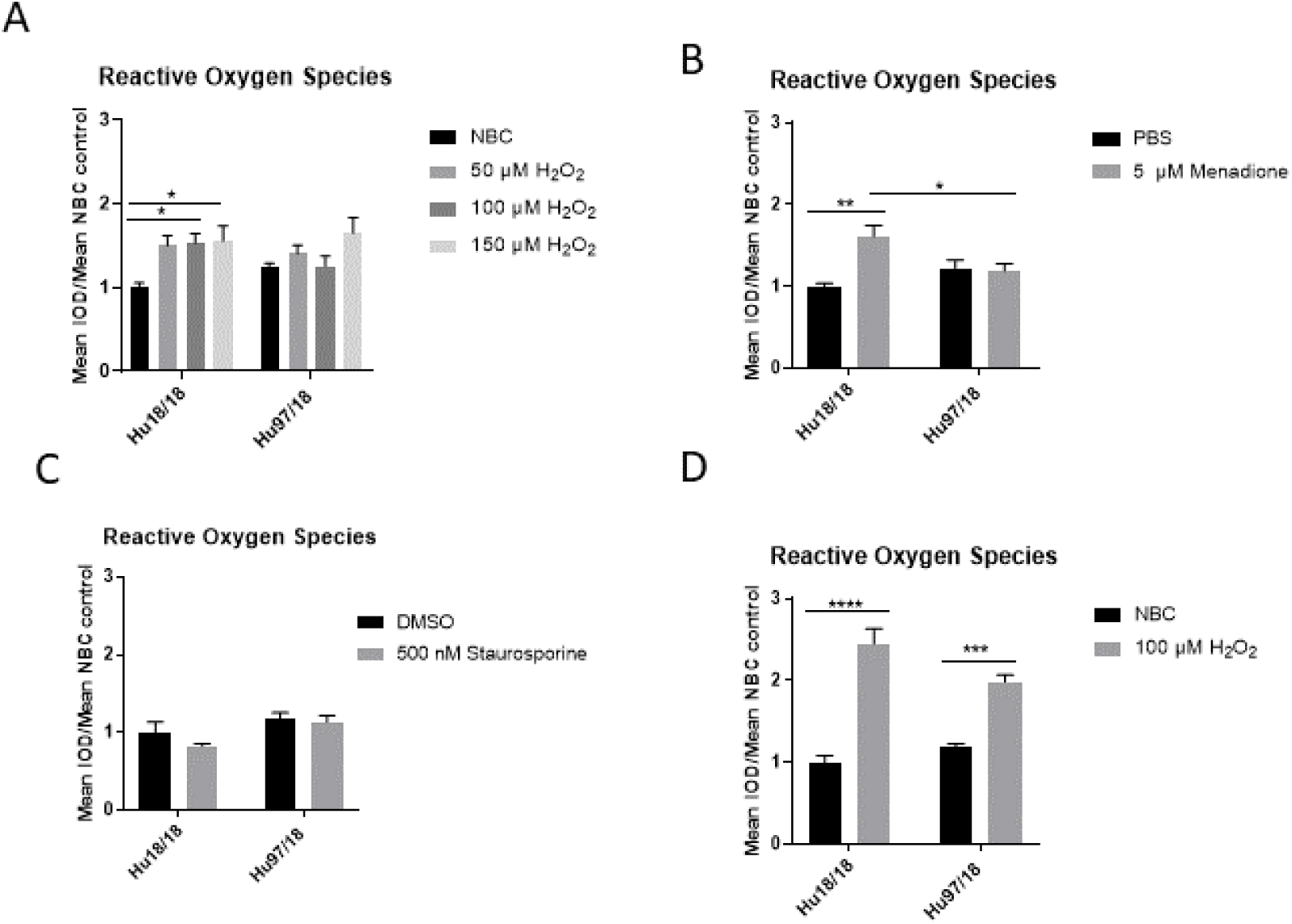
Oxidative stressors increase ROS in immature control and mature HD primary neurons. Primary Hu97/18 and Hu18/18 control neurons were treated with stressors for 24h and ROS was measured by live-imaging of CM-H2DCFDA florescence. Immature neurons were treated with (**A**) NBC or H_2_O_2_ at the indicated doses. (**B**) PBS or 5µM menadione. (**C**) DMSO or 500nM staurosporine. (**D**) Mature neurons were treated with 100µM H_2_O_2_. All data normalized to mean Hu18/18 vehicle control. *=difference between indicated bars, *p<0.05, **p<0.01, ***p<0.001, ****=p<0.0001. Error bars ± SEM.

These initial experiments were conducted at day *in vitro* (DIV)13 for consistency with previously published studies of the effects of stressors in primary neurons (Graham et al., 2009; Tsvetkov et al., 2010; Zeron et al., 2002). However, primary neurons do not begin to reach full maturity and become maximally electrically active until DIV21 (Biffi, Regalia, Menegon, Ferrigno, & Pedrocchi, 2013). Because of this, we repeated our initial experiments in neurons at DIV21. Contrary to what we saw in immature HD neurons, mature HD neurons exhibit the expected increase in ROS upon H_2_O_2_ treatment **(Figure 1D**, 2 way ANOVA treatment p<0.0001, genotype p=0.2505, post-test vs. NBC same genotype Hu18/18: p<0.0001; Hu97/18: p=0.0003). This demonstrates that mature neurons are a more relevant model of cellular function and dysfunction in adult onset diseases.

### Mature HD neurons have elevated basal oxidative damage that is not exacerbated by H_2_O_2_ treatment

ROS are naturally detoxified by endogenous antioxidants in the cell (Pham-Huy, He, & Pham-Huy, 2008). However, when high levels of ROS flood the cell, they can overwhelm antioxidants and cause damage to DNA, RNA, and proteins (Schieber & Chandel, 2014). Increased DNA damage has been reported in HD (Bogdanov, Andreassen, Dedeoglu, Ferrante, & Beal, 2001; Brocardo, McGinnis, Christie, & Gil-Mohapel, 2016; Susan E. Browne, Ferrante, & Beal, 1999; C.-M. Chen et al., 2007; Polidori, Mecocci, Browne, Senin, & Beal, 1999; Sorolla et al., 2008). The antioxidant and DNA damage response has also been reported to be diminished in HD neurons (Beal et al., 1992; C.-M. Chen et al., 2007; Peña-Sánchez et al., 2015). Therefore, we interrogated whether HD neurons are hypersensitive to oxidative stress by quantifying 8-Oxo-2’-deoxyguanosine (8-Oxo-dG), a marker of DNA oxidation. We found an increase in oxidative damage in immature control, but not HD neurons when treated with H_2_O_2_ **(Figure 2A,B**, 2 way ANOVA treatment p=0.0420, genotype p=0.0101, post-test vs. NBC same genotype Hu18/18: 50µM p=0.3024, 100µM p<0.0001; Hu97/18: 50µM p=0.1877, 100µM p=0.1467). Because oxidative damage becomes significantly elevated in Hu18/18 neurons at 100 µM, this dose was chosen for subsequent experiments. Interestingly, we observed similar effects in mature neurons; oxidative damage was induced in mature control, but not HD neurons (**Figure 2C,D**, 2 way ANOVA treatment p=0.0140, genotype p=0.1973, post-test vs. NBC same genotype Hu18/18 p=0.0140; Hu97/18 p=0.5905). However, we did observe elevated basal levels of oxidative DNA damage in mature, but not immature HD neurons (**Figures 2B,D**, post-test vs. NBC Hu18/18 same DIV, immature p=0.3186, mature p=0.0010), resulting in damage similar to that observed in H_2_O_2_ treated control neurons. This is consistent with previous findings demonstrating elevated oxidative damage in HD neurons (S. E. Browne et al., 1997; Polyzos et al., 2016), further supporting the use of mature neurons in mechanistic studies.

**Figure 2.**
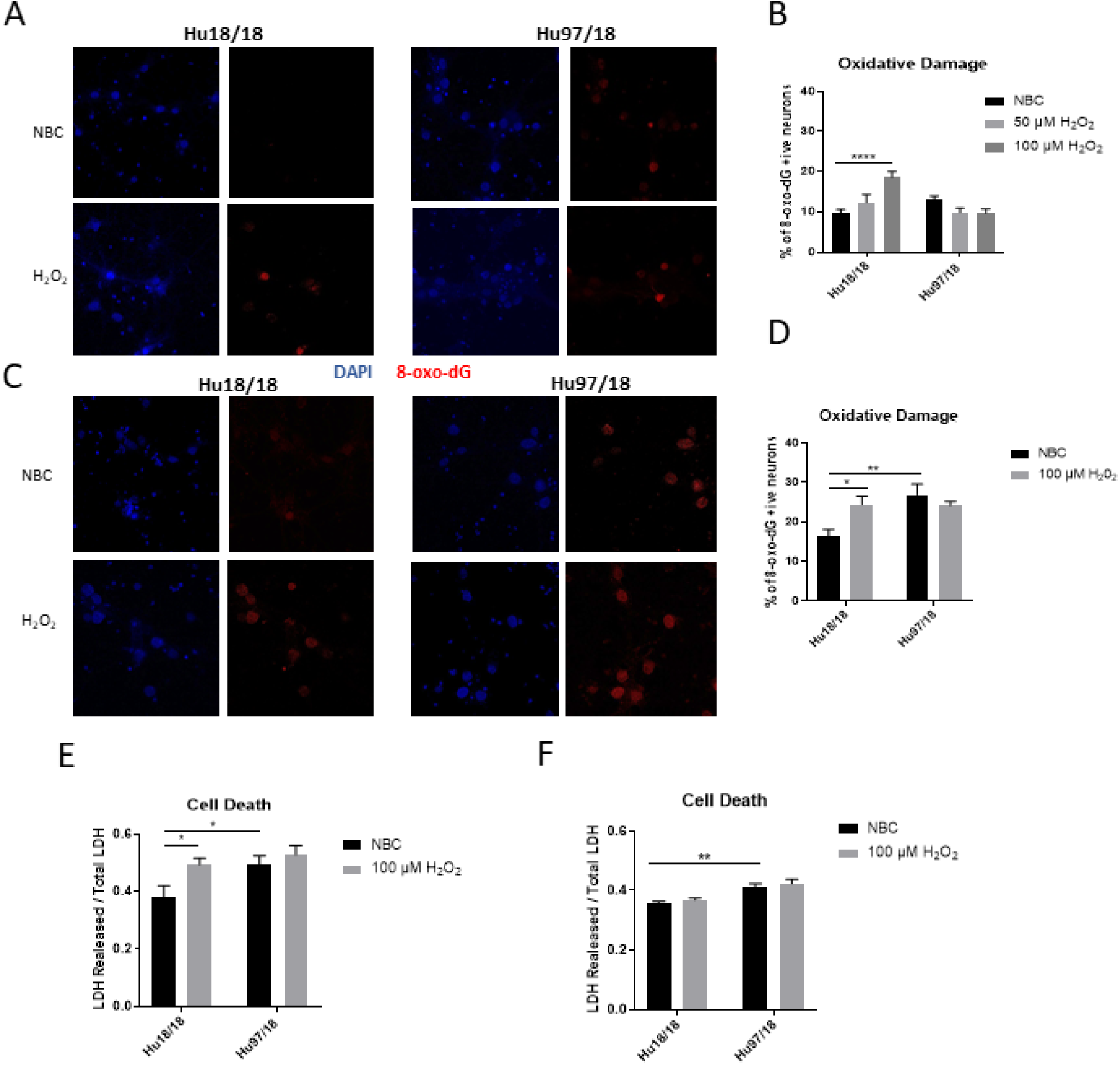
Oxidative stress does not induce oxidative damage or cell death in HD primary neurons. Primary Hu18/18 and Hu97/18 neurons were treated with NBC vehicle or 100µM H_2_O_2_ for 24h. (**A-D**) DNA damage was assessed by 8-Oxo-dG immunocytochemistry. (**A,B**) Representative images (**A**) and quantification (**B**) of % 8-Oxo-dG positive immature neurons. (**C,D**) Representative images (**C**) and quantification (**D**) of % 8-Oxo-dG positive mature neurons. (**E,F**) Cell death was assessed by quantifying LDH released/total in (**E**) immature and (**F**) mature neurons. *=difference between indicated bars, *p<0.05, **p<0.01, ****=p<0.0001. Error bars ± SEM.

To determine whether the absence of observed H_2_O_2_-induced DNA damage in HD neurons resulted solely from increased cell death, we measured lactate dehydrogenase (LDH) release in basal and H_2_O_2_ treated conditions. We found H_2_O_2_ treatment caused significant cell death in immature control, but not HD neurons (Figure 2E treatment p=0.0220, genotype p=0.0213, post-test vs. NBC same genotype Hu18/18: p=0.0270; Hu97/18: p=0.8742). However, basal levels of cell death were increased in immature HD neurons (**Figure 2E**, post-test vs NBC Hu97/18 p=0.0345). In mature neurons, H_2_O_2_ did not induce cell death in control or HD neurons. However HD neurons displayed increased levels of basal cell death (**Figure 2F**, treatment p=0.2289, genotype p<0.0001, post-test Hu97/18 NBC vs. Hu18/18 NBC p=0.0011). Together, these data demonstrate that mature HD neurons have some mild phenotypes which are not exacerbated by exogenous oxidative stress.

### H_2_O_2_ treatment does not induce canonical stress response pathways in HD neurons

Caspase-3 is a protease that, when activated, initiates the cellular apoptosis cascade, which can be the ultimate result of oxidative stress. Wild-type HTT (wtHTT) inhibits caspase-3 activation, and such inhibition attenuates neurodegeneration in HD mouse brain (Zhang et al., 2006). To determine if caspase-3 signaling is altered in HD neurons or in response to H_2_O_2_ treatment, we measured active caspase-3 protein. We found that in immature neurons, caspase-3 was not induced in control or HD neurons after H_2_O_2_ treatment. However, in mature control neurons, we saw a strong trend toward increased activation of caspase-3 post-H_2_O_2_ treatment, suggesting a potential induction of caspase-3 activation. HD neurons did not exhibit this trend (**Figure 3A-D**, Immature neurons: treatment p=0.2988, genotype p=0.3600, post-test vs NBC same genotype Hu18/18 p>0.9999, Hu97/18 p=0.6149; mature neurons: treatment p=0.2988, genotype p=0.36, post-test vs NBC same genotype Hu18/18 p=0.0603, Hu97/18 p>0.9999). This suggests that HD neurons do not mount the apoptotic cascade in response to oxidative stress. This may be either because damage is not sufficient to activate caspase-3 or because HD neurons have a deficiency in this stress response. This also suggests that the elevated basal cell death we observed in Hu97/18 neurons may not be due to caspase-3-dependent apoptosis.

**Figure 3.**
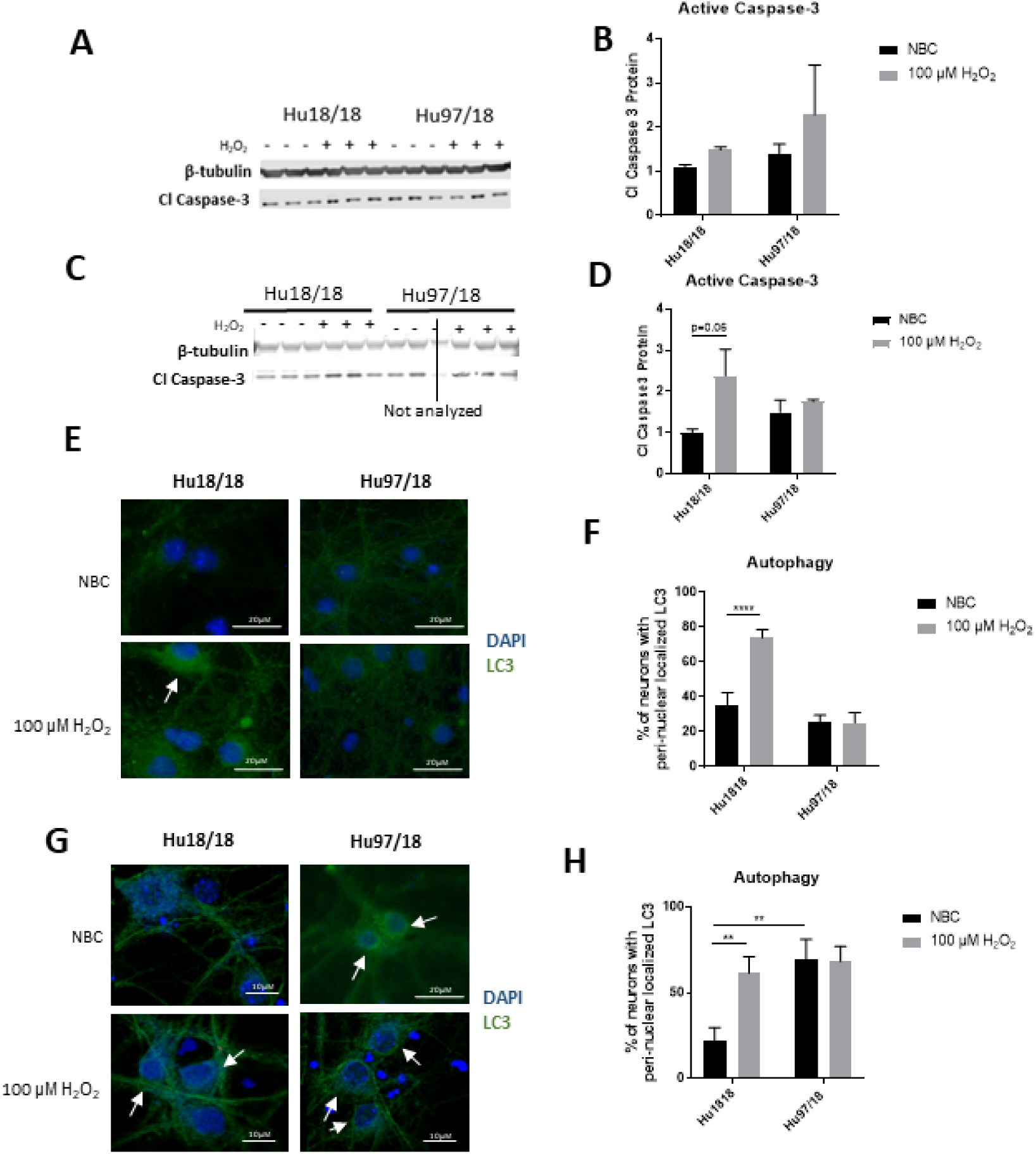
H_2_O_2_ induces autophagic and possibly caspase-3 apoptotic pathways in control but not HD neurons. Primary Hu18/18 and Hu97/18 neurons were treated with NBC vehicle or 100µM H_2_O_2_ for 24h. (**A-D**) Apoptosis was assessed by quantifying active caspase-3 protein by WB. (**A,B**) representative images (**A**) and quantification (**B**) in immature neurons. (**C,D**) Representative images (**C**) and quantification (D) in mature neurons. (**E-G**) Autophagy was assessed by LC3 immunocytochemistry. (**E,F**) representative images (**E**) and quantification (**F**) of % of immature neurons with perinuclear localized LC3. (**G,H**) Representative images (**G**) and quantification (**H**) of % of mature neurons with perinuclear localized LC3. *=difference between indicated bars, **p<0.01, ****=p<0.0001. Error bars ± SEM.

H_2_O_2_ has previously been shown to induce autophagy, which decreases damage in cells and protects cells from cell death at low levels, and, at higher levels, can induce autophagy-dependent apoptosis (L. Chen, Liu, Yin, Luo, & Huang, 2009; Higgins, Devenish, Beart, & Nagley, 2011). In fact, a previous study showed that cell death in response to H_2_O_2_ treatment is mediated through autophagic activity in cultured primary cortical neurons (L. Chen et al., 2009; Higgins et al., 2011). However, autophagy is impaired in HD, with HD cells exhibiting cargo recognition and loading defects that result in accumulation of empty autophagosomes (Croce & Yamamoto, 2019; Martin, Ladha, Ehrnhoefer, & Hayden, 2015; M. Martinez-Vicente et al., 2010). To investigate the autophagic response in HD primary neurons, we treated control and HD neurons with H_2_O_2_ and assessed LC3 localization. LC3-positive vesicles can be found in both the soma and axons of a neuron, but undergo retrograde transport to localize with lysosomes in a perinuclear fashion under stress (S. Lee, Sato, & Nixon, 2011). We found increased perinuclear LC3 in response to H_2_O_2_ treatment in immature control, but not HD neurons (**Figure 3E,F**, treatment p=0.0016, genotype p<0.0001, post-test vs. NBC same genotype Hu18/18 p<0.0001, Hu97/18 p>0.9999). Similarly, in mature neurons, we observed induction of perinuclear LC3 localization in control neurons, and no effect in HD neurons (**Figure 3G,H**, treatment p=0.0460, genotype p=0.0057, post-test vs. NBC same genotype Hu18/18 p=0.0056, Hu97/18 p>0.9999). However, we also saw increased basal perinuclear LC3 in mature HD neurons (Hu97/18 NBC vs. Hu18/18 NBC p=0.0018), demonstrating dysregulated autophagy only in mature HD neurons.

### Progerin treatment induces aging and uncovers stress-related phenotypes in HD primary neurons

Primary neurons, even when matured in culture, are still developmental, with no markers of aging. Because HD is a disease of aging, developmental neurons may not be the best model. In an effort to provide a more relevant neuronal HD model, we used progerin treatment to induce aging. Progerin is a truncated form of the Lamin A protein that causes Hutchinson-Gilford Progeria Syndrome, a disease of premature aging (Eriksson et al., 2003). Progerin expression causes cellular senescence in normal fibroblasts, and accumulation also occurs during normal aging (Cao et al., 2011; McClintock et al., 2007). Progerin expression has previously been used to induce aging and uncover aging-related phenotypes in iPSC-derived neurons from Parkinson disease patients (Miller et al., 2013).

To induce aging in mature neurons, we used an adeno-associated virus (AAV2/1-GFP-progerin) or nuclear GFP control (AAV2/1-nGFP). Because the previous study in iPSCs used a 5 day progerin treatment interval, and in previous work we have observed maximal transgene expression ∼4 days post-transduction with this serotype in cultured cells (Southwell, Ko, & Patterson, 2009), we selected a 9 day post-transduction interval. In preliminary experiments using AAV2/1-GFP in differentiating iPSC colonies, we observed efficient transduction and high transgene expression after this interval only in differentiating cells, and not in undifferentiated iPSCs (**Figure S2A**). In a dosing study in striatal-like neurons derived from iPSCs, we found efficient transduction at e10 viral genomes/ml (VG/ml), so chose this dose for our experiments (**Figure S2B**).

We first investigated whether progerin could induce markers of age in primary cortical neurons. We found that progerin treatment caused nuclear blebbing, which is a sign of cellular senescence, in neurons irrespective of genotype (**Figure 4A**). We also quantified proteins known to change with age. NMDAR1 is a subunit of NMDA receptors that decreases throughout aging (Gazzaley, Weiland, McEwen, & Morrison, 1996). We found that inducing aging with progerin caused a trend toward decreased NMDAR1 levels in control but not HD neurons (**Figure 4B**, treatment p=0.2436, genotype p=0.7193, post-test vs. nGFP same genotype Hu18/18 p=0.1341, Hu97/18 p>0.9999). Dynamin 1 helps facilitate endocytosis in neurons, and has been found to decrease with both aging and in neurodegenerative disease (Yoo et al., 2016). We found a trend toward a decrease in Dynamin 1 with progerin treatment in control neurons that we didn’t find in HD neurons (**Figure 4C**, treatment p=0.4405, genotype p=0.1068, post-test vs. nGFP same genotype Hu18/18 p=0.0611, Hu97/18 p=0.3572). Lamin B1 is a nuclear lamin protein that has previously been shown to decrease with age (Freund, Laberge, Demaria, & Campisi, 2012)(Wang, Ong, Chojnowski, Clavel, & Dreesen, 2017). We saw a dramatic decrease in Lamin B1 protein in progerin-treated control neurons (**Figure 4D**, treatment p=0.0183, genotype p=0.1711, post-test vs. nGFP same genotype Hu18/18 p=0.0129, Hu97/18 p>0.9999). Interestingly, we also saw that HD neurons basally had a trend toward decreased NMDAR1 (**Figure 4B**, Hu97/18 nGFP vs. Hu18/18 nGFP p=0.3485), a significant decrease in dynamin 1 (**Figure 4C**, Hu97/18 nGFP vs. Hu18/18 nGFP p= p=0.03572), and a strong trend toward decreased Lamin B1 (**Figure 4D**, Hu97/18 nGFP vs. Hu18/18 nGFP p= p=0.0605). These results are consistent with other reports of advanced biological age in HD brain and neurons (Grima et al., 2017; Horvath et al., 2016). It is also important to note that in mature neurons, treatment with progerin caused cellular rearrangement of HD neurons (**Figure S3**), which has previously been observed in Hu97/18 primary neurons treated with poorly tolerated antisense oligonucleotides (Skotte et al., 2014). Together, this demonstrates that progerin treatment alters levels of aging markers and that mature HD neurons display some accelerated markers of aging.

**Figure 4.**
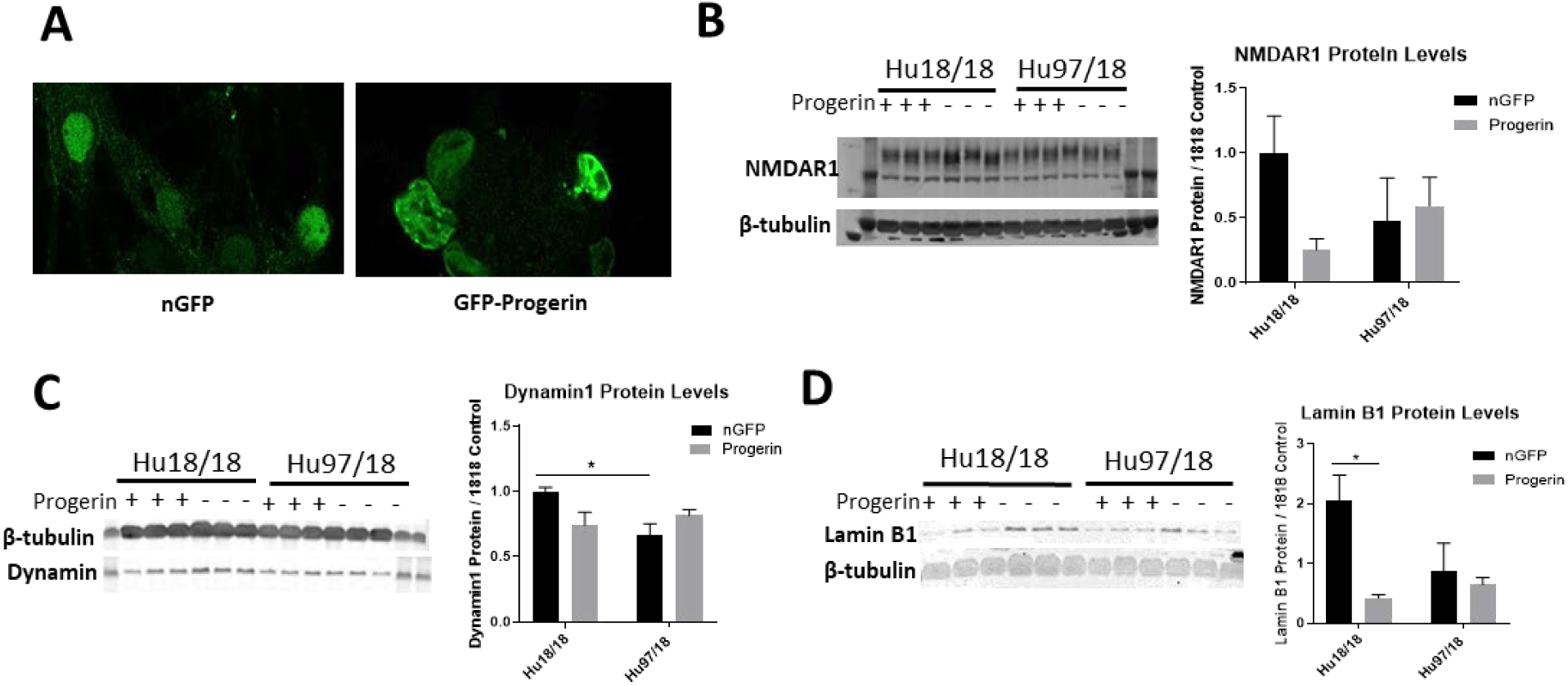
Progerin treatment induces aging in primary neurons. Mature primary Hu18/18 and Hu97/18 neurons were treated with nGFP or progerin. (**A**) Progerin induces nuclear blebbing, which occurs with age. (**B-D**) WB analysis of proteins known to decrease over natural aging; (**B**) NMDAR1, (**C**) Dynamin1, and (**D**) Lamin B1. *=difference between indicated bars, *p<0.05. Error bars ± SEM.

mtHTT oligomerization and aggregation is a characteristic of HD brains and neurons, but something that is not observed in embryonic HD primary neurons. We assessed mtHTT aggregation by immunoreactivity of EM48, an anti-HTT antibody that recognizes only aggregated or oligomeric mtHTT. Mature HD primary cortical neurons entirely lack EM48 immunoreactivity, suggesting that mtHTT is primarily in a monomeric state within these cells. However, after progerin-induced aging, we observed EM48 immunoreactivity (**Figure 5A**), suggesting that mtHTT has undergone some form of aggregate seeding and/or oligomerization. Thus, that biological age can modulate the folding and aggregation state of mtHTT protein irrespective of the passage of time. This is consistent with a previous report of EM48 positive mtHTT aggregates in striatal-like neurons directly converted from HD patient fibroblasts, which maintain fibroblast epigenetics, including markers of age (Victor et al., 2018).

**Figure 5.**
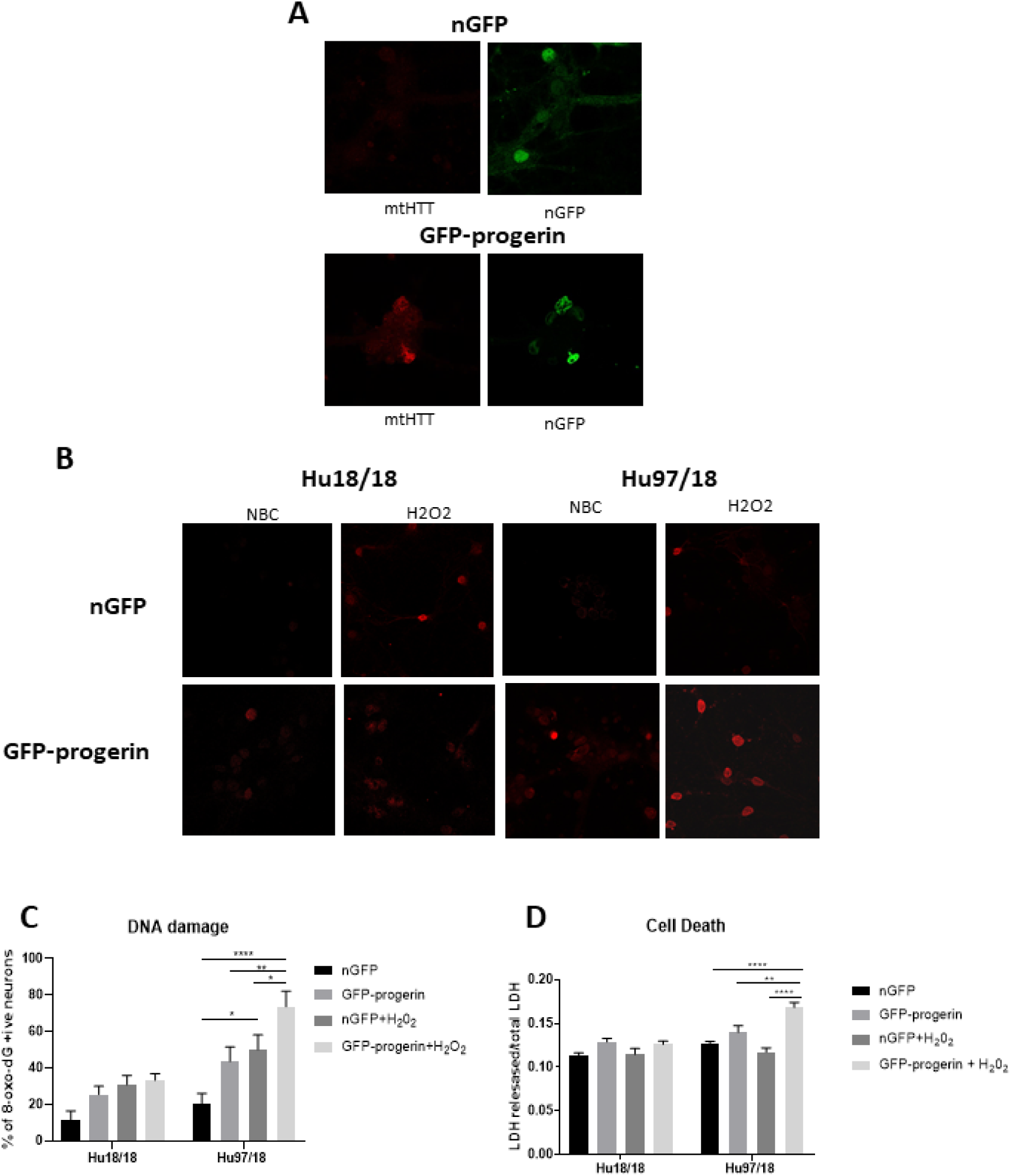
Progerin induced aging causes aggregate seeding/oligomerization of mtHTT and increased susceptibility to oxidative stress induced DNA damage and cell death in HD neurons. (**A**) Mature Hu18/18 and Hu97/18 neurons were treated with nGFP or progerin to induce aging, and mtHTT was assessed by EM48 immunocytochemistry. (**B,C**) Mature Hu18/18 and Hu97/18 neurons were aged with progerin or given nGFP control followed by H_2_O_2_ or vehicle control treatment. Oxidative DNA damage was assessed by oxo-8-dG immunocytochemistry. Representative images (**B**) and quantitation of % of neurons exhibiting DNA damage (**C**).(**D**) Mature primary Hu18/18 and Hu97/18 neurons were treated with nGFP or progerin followed by H_2_O_2_ or vehicle control. Cell death was assessed by quantifying released/total LDH. *=difference between indicated bars, *=p<0.05, **p<0.01, ****=p<0.0001. Error bars ± SEM.

We next investigated how accelerated aging affects sensitivity to oxidative stress by treating neurons with nGFP (nGFP+vehicle), progerin (GFP-progerin+vehicle), H_2_O_2_ (nGFP+ H_2_O_2_) or the combination (progerin+ H_2_O_2_). We found a strong trend toward a selective increase in oxidative DNA damage in HD, but not control, neurons when aged with progerin (**Figure 5B,C, S4-S5**, treatment p<0.0001, genotype p<0.0001, post-test vs. nGFP same genotype Hu18/18 progerin p=0.8699; Hu97/18 progerin p=0.1088. Similar results were seen for cells treated with H_2_O_2_ (post-test vs. nGFP same genotype Hu18/18 H_2_O_2_ p=0.1919; Hu97/18 H_2_O_2_ p=0.0131). Interestingly, we found the combination of aging and oxidative stress to synergistically increase oxidative damage selectively in HD neurons beyond what was seen with either insult alone (post-test vs. progerin+H_2_O_2_ same genotype Hu18/18 nGFP p=0.0733, progerin p>0.9999, H_2_O_2_ p>0.9999; Hu97/18 nGFP p<0.0001; progerin p=0.0069, H_2_O_2_ p=0.0500). Additionally, we found that the combination of progerin-induced aging and H_2_O_2_-mediated oxidative stress had no effect on cell death in control neurons. Conversely, in HD neurons, this combination did induce cell death where either insult alone did not (**Figure 5D**, treatment p<0.0001, genotype p<0.0001, post-test vs. nGFP same genotype Hu18/18 progerin p=0.1679, H_2_O_2_ p>0.9999, progerin+ H_2_O_2_ p=0.3909; Hu97/18 progerin p=0.3781, H_2_O_2_ p>0.9999, progerin+ H_2_O_2_ p<0.0001). Together these results demonstrate that progerin-induced aging uncovers HD-related phenotypes, and that there is a synergistic effect of aging and cellular stress on dysfunction and disease of HD neurons.

### Inducing aging with progerin uncovers HD phenotypes in iPSC-derived human neurons

iPSC-derived neurons from patients can be a powerful *in vitro* tool to study disease processes and mechanisms. However, iPSC-derived neurons from patients with diseases of aging have few robust phenotypes (Wu, Chiu, Yeh, & Kuo, 2019). Adult-onset HD iPSC-derived neurons have very mild phenotypes and often require the addition of stress (HD iPSC Consortium, 2012). To investigate if inducing aging with progerin could uncover phenotypes in HD iPSC-derived neurons, we infected HD and control iPSC-derived striatal-like neurons with AAV2/1-GFP-progerin or AAV2/1-nGFP control. Similar to primary neurons, we found that progerin-induced aging causes nuclear blebbing in neurons irrespective of genotype (**Figure 6A**). Dendritic length and complexity are two measures of neuronal health that decrease with age and throughout the course of HD (Buren, Tu, Parsons, Sepers, & Raymond, 2016; Lerner, Trejo, Zhu, Chesselet, & Hickey, 2012; Schmidt et al., 2018). We found that inducing aging with progerin decreases dendritic length and radial dendritic complexity of neurons (**Figure 6B-D**, treatment p=0.0006, genotype p=0.5275, post-test vs. nGFP same genotype Hu18/18 p=0.0294, Hu97/18 p=0.0280). This could be indicative of reduced systems level connectivity. Similar to primary neurons, we observed a trend toward caspase 3 activation in response to progerin-induced aging in neurons (**Figure 6E**, treatment p=0.1060, genotype p=0.4132, post-test vs. nGFP same genotype Hu18/18 p=0.4182, Hu97/18 p=0.6036), indicating that our aging paradigm itself does not induce the apoptotic cascade.

**Figure 6.**
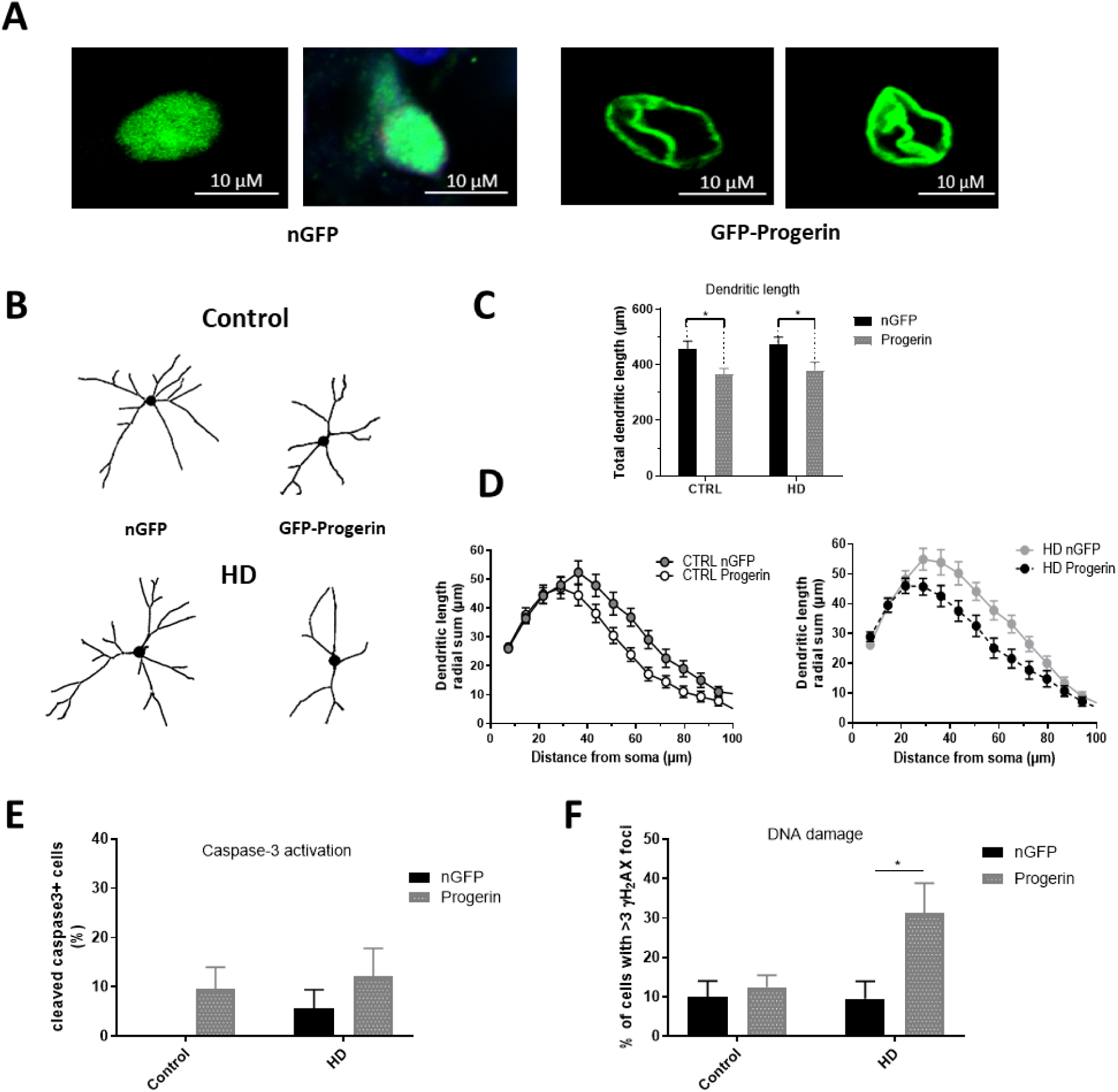
Progerin treatment induces aging and unmasks HD-like phenotypes in HD iPSC-derived neurons. HD patient and healthy control iPSCs were differentiated toward striatal-like neurons and treated with nGFP or progerin and evaluated by immunocytochemistry. (**A**) GFP staining demonstrates nuclear blebbing in progerin, but not nGFP treated neurons. (**B-D**) Dendritic morphology was assessed in transduced cells by (**C**) total dendrite length and (**D**) radial dendritic complexity. (**E**) Apoptosis was assessed by % of transduced neurons that are caspase-3 positive. (**F**) DNA damage was assessed by % of transduced neurons with >3 γH_2_AX foci, revealing a selective increase in HD neurons. *=difference between indicated bars. *p<0.05. Error bars ± SEM.

We also assessed DNA damage in neurons by immunocytochemistry of γH_2_AX foci, and observed selectively increased DNA damage in aged HD neurons (**Figure 6F**, treatment p=0.0514, genotype p=0.1383, post-test vs. nGFP same genotype Hu18/18 p>0.9999, Hu97/18 p=0.0190). This demonstrates unmasking of an HD-like phenotype by accelerated biological age irrespective of the passage of time, further supporting a role for aging in the pathogenesis of HD.

### Progerin treatment induces aging and oxidative damage in the brain of HD mice

We next wanted to evaluate if expression of progerin in the brain of HD mice could induce aging-related phenotypes as was observed in primary and iPSC-derived neurons. The YAC128 mouse model harbors a full-length human mutant *HTT* transgene and recapitulates many of the behavioral and neuropathological features of HD (Carroll et al., 2011; Slow et al., 2003; Southwell et al., 2009). YAC128 mice received bilateral instriatal infusions by convection-enhanced delivery of either AAV2/1-GFP-progerin or AAV2/1-GFP delivered at a dose of 1e10 VG/hemisphere at 2 months of age and were collected 8 weeks post-injection for either immunohistochemistry (IHC) or western blot (WB) analysis.

Broad distribution throughout the striatum and deeper layers of the cortex was observed with both AAV2/1-GFP (**Figure S6A**) and AAV2/1-GFP-progerin (**Figure S6B**). Consistent with the reported tropism of this AAV serotype, we observed efficient transduction of MSNs (**Figure S7A**), neurons (**Figure S7B**, as well as astrocytes and neural progenitors (**Figure S7C**) in the striatum. Similar to our culture models, nuclear blebbing was observed in cells expressing GFP-progerin (**Figure 7A**).

**Figure 7.**
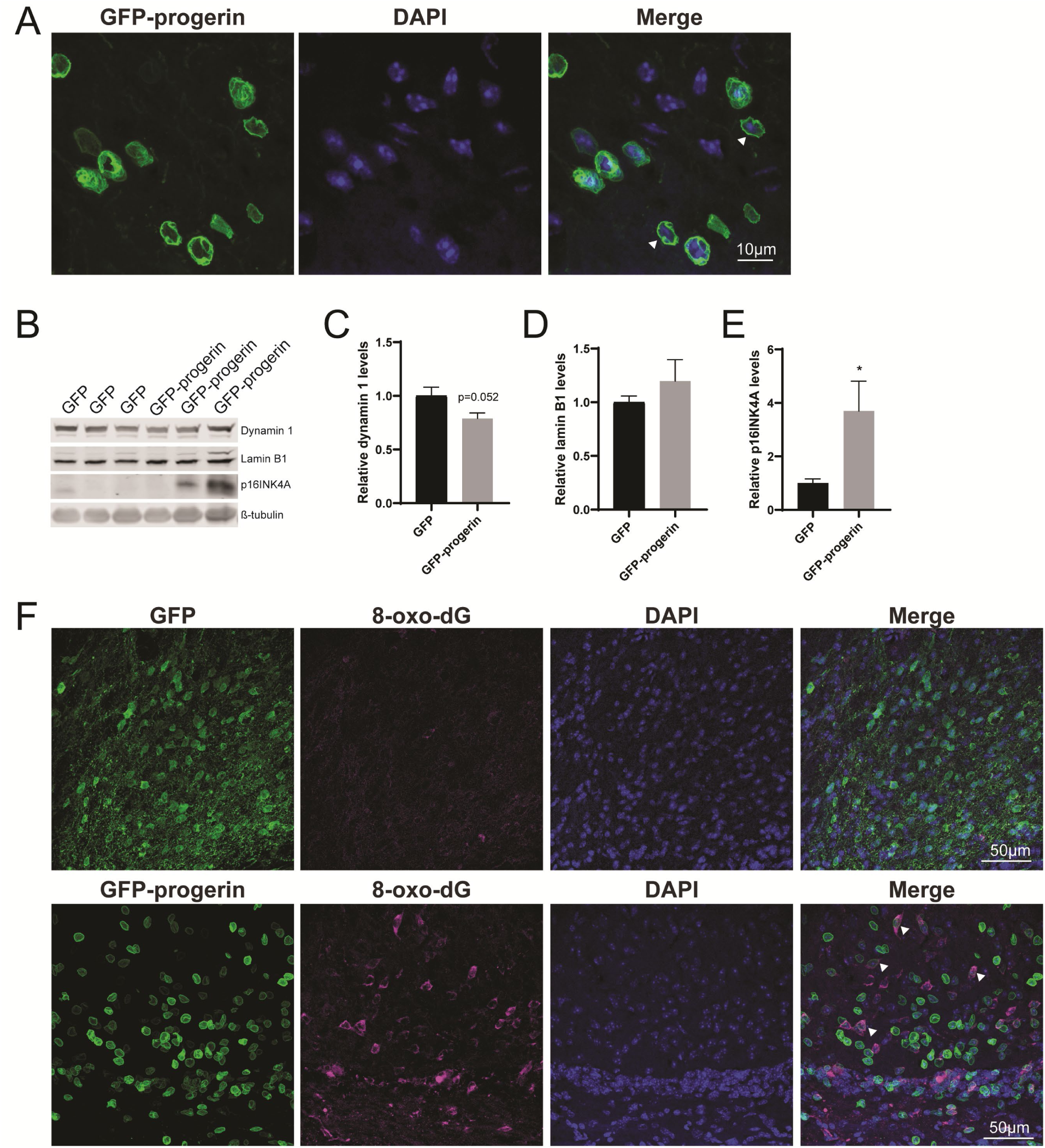
Progerin treatment induces aging and DNA damage in the HD mouse brain. **(A)** Representative confocal image of YAC128 brain treated with AAV2/1-GFP-progerin shows nuclear blebbing. Scale bar = 10µm. Representative Western blots **(B)** and quantifications of aging markers **(C)** dynamin 1, **(D)** lamin B1 and **(E)** p16INK4a. Error bars ± SEM. *=difference from GFP. *p<0.05 **(F)** Confocal images of 8-oxo-dG staining in the YAC128 mouse brain treated with either GFP or GFP-progerin.

To evaluate the effect of progerin on markers of aging in YAC128 mice, striatal tissues from GFP and GFP-progerin treated mice were assessed for dynamin 1, lamin B1 and p16INK4a by WB (**Figure 7B**). We observed a strong trend towards a decrease in dynamin 1 levels in YAC128 mice in response to progerin-induced aging but this did not reach significance (**Figure 7B,C**, Unpaired t-test p=0.052). We did not observe any effect of progerin treatment on lamin B1 levels (**Figure 7B, D**, Unpaired t-test p=0.3630). p16INK4a is cyclin-dependent kinase inhibitor involved in the regulation of the cell cycle which has been found to be a robust *in vivo* marker of cellular aging (Ressler et al., 2006). We observed a significant increase in p16INK4a levels with progerin treatment (**Figure 7B, E**, Unpaired t-test *p=0.0373). These data suggest that progerin can induce selected markers of aging in the brain of YAC128 mice. It is important to note that lysates represent an average of all the cells in that tissue, and that the effects of progerin treatment on levels of aging makers in these striatal lysates may be blunted as a result of cells that were not transduced by AAV2/1-GFP-progerin.

We also qualitatively assessed the effect of progerin on oxidative damage in YAC128 mice by staining for 8-oxo-dG in brains treated with either GFP or GFP-progerin. We observed minimal staining for 8-oxo-dG in the striatum of YAC128 mice treated with AAV2/1-GFP. Conversely, we observed strong staining for 8-oxo-dG in the striatum and cortex of AAV2/1-GFP-progerin treated YAC128 mice. There was also a high degree of overlap between 8-oxo-dG and progerin transduced cells. This data suggests that progerin treatment induces oxidative DNA damage in the brain of HD mice.

## Discussion

Oxidative stress and damage have been observed in many models of HD and in post-mortem brains of HD patients (Bogdanov et al., 2001; S. E. Browne et al., 1997; C.-M. Chen et al., 2007; Johri & Beal, 2012a; Klepac et al., 2007; Polidori et al., 1999; Sepers & Raymond, 2014). Oxidative stress can cause not only the damage of organelles and apoptosis, but also can play a role in somatic expansion of the CAG repeat in HD (Jonson, Ougland, Klungland, & Larsen, 2013). HTT is a critical component of multiple cellular stress responses (Atwal et al., 2007; Munsie et al., 2011; Nath, Munsie, & Truant, 2015) In response to various stressors, including oxidative stress, HTT reversibly localizes with early endosomes and forms distinct cytosolic puncta (Nath, Munsie, & Truant, 2015). Cells expressing mtHTT are defective at recovering from this response effectively (Nath et al., 2015). Additionally, HTT can act as a scaffold protein of the DNA repair pathway in response to oxidative stress, and that this response is impaired in the presence of mtHTT (Maiuri et al., 2017). Together, these provide evidence for mtHTT directly causing oxidative damage in cells by diminishing proper stress responses. However, whether oxidative damage is a driver of disease pathogenesis has been debated.

Here, we demonstrate differences in the oxidative stress response between immature and mature primary cortical neurons from HD mice. We observe decreased susceptibility to oxidative stress in immature HD neurons compared to control or mature HD neurons. This is an important distinction because several experiments investigating the role of stress in HD in primary cortical neurons have been performed in neurons at DIV13 (Graham et al., 2009; Tsvetkov et al., 2010; Zeron et al., 2002), which could potentially skew conclusions drawn from these studies. We found that while Hu97/18 immature neurons had slightly elevated basal levels of ROS, treatment with oxidative stressors did not induce ROS or oxidative damage in neurons. This could potentially be due to optimized endogenous antioxidant mechanisms to clear ROS in immature Hu97/18 neurons that decline with neuronal maturity, though we find this unlikely considering that cellular stress responses are impaired in HD neurons.

We did not observe cell death in either control or HD neurons following 100µM H_2_O_2_ treatment, which is consistent with previous findings that treatment with H_2_O_2_ up to 500µM does not increase LDH release in control or HD (ST*Hdh*^Q111^) neurons (Jin, Hwang, Jo, & Johnson, 2012). However, we did find increased cell death at a basal level in mature HD neurons. Although this, to our knowledge, has not been previously reported in HD primary cortical neurons, viral expression of HTT fragments with expanded CAG in neurons causes increased cell death over control HTT expression (Grima et al., 2017; Hermel et al., 2004). Additionally, several groups have reported increased caspase activation and cytochrome c release in HD models, suggestive of activated apoptotic cascades in HD neurons (Dickey et al., 2017; Hermel et al., 2004; Kim et al., 2010; Li, Lam, Cheng, & Li, 2000; Yang et al., 2010). We found a trend toward increased caspase-3 activation in Hu97/18 neurons, and while this didn’t reach significance, we do believe this pathway could be contributing to cell death in these neurons. Additionally, we observed elevated LC3 perinuclear localization indicative of increased autophagy in HD neurons, which at high levels, can lead to apoptosis (Gump & Thorburn, 2011) or to accumulation of empty autophagasomes due to cargo recognition deficits as have been reported in HD mouse brain and patient lymphoblasts (Marta Martinez-Vicente et al., 2010). Both of which could be in accordance with previous reports of increased LC3 expression HD neurons (Ehrnhoefer et al., 2018; Lee et al., 2015).

In addition to basal differences between immature and mature neurons, we also found that progerin treatment not only effectively induces aging in HD mouse primary cortical neurons, HD patient iPSC-derived neurons and the brains of HD mice, it also induces mtHTT oligomerization, selective DNA damage, and sensitized neurons to oxidative stress. Most cases of HD are adult-onset; however, most *in vitro* models of HD are embryonic-like and even *in vivo* models are young at phenotypic onset, such as the R6/2 mouse, which has previously been found to have increased DNA damage compared to controls independent of age (Goula et al., 2009).

The evidence here, along with previous work showing progerin-induced aging can uncover phenotypes in iPSC-derived neurons from Parkinson disease patients (Miller et al., 2013), provides rationale for using progerin-induced aging as a paradigm to better model HD and other age-dependent neurodegenerative diseases. This will be especially important for HD, as neuronal and mouse models harboring the most common pathogenic (adult-onset) CAG repeat lengths show very few phenotypes (Consortium, 2012; Hodgson et al., 1999).

The obstacle of modeling HD in developmental neurons has been addressed by another group through direct conversion of HD patient fibroblasts to striatal-like neurons (Victor et al., 2018). Directly converted neurons maintain the donor fibroblast epigenetic characteristics, including those of age and, like progerin-aged neurons, display HD-like phenotypes not seen in developmental HD or control neurons. The consistent results between these aged human neuron models; directly converted neurons with the epigenetics of skin cells and iPSC-derived neurons after progerin-induced aging, suggests that this is a true feature of HD neurons rather than a model system artifact. Additionally, the consistent elevation of DNA damage selectively in HD neurons across the three model systems employed in this study provides strong evidence that age-related changes in stress and damage responses may be primary drivers of HD pathogenesis. This is particularly important because DNA damage may drive somatic expansion in HD neurons, which, in turn, may drive HD onset and progression, (Dragileva et al., 2009; Swami et al., 2009; Wheeler et al., 2003), as shown in our proposed model of HD pathogenesis (**Figure 8**).

**Figure 8.**
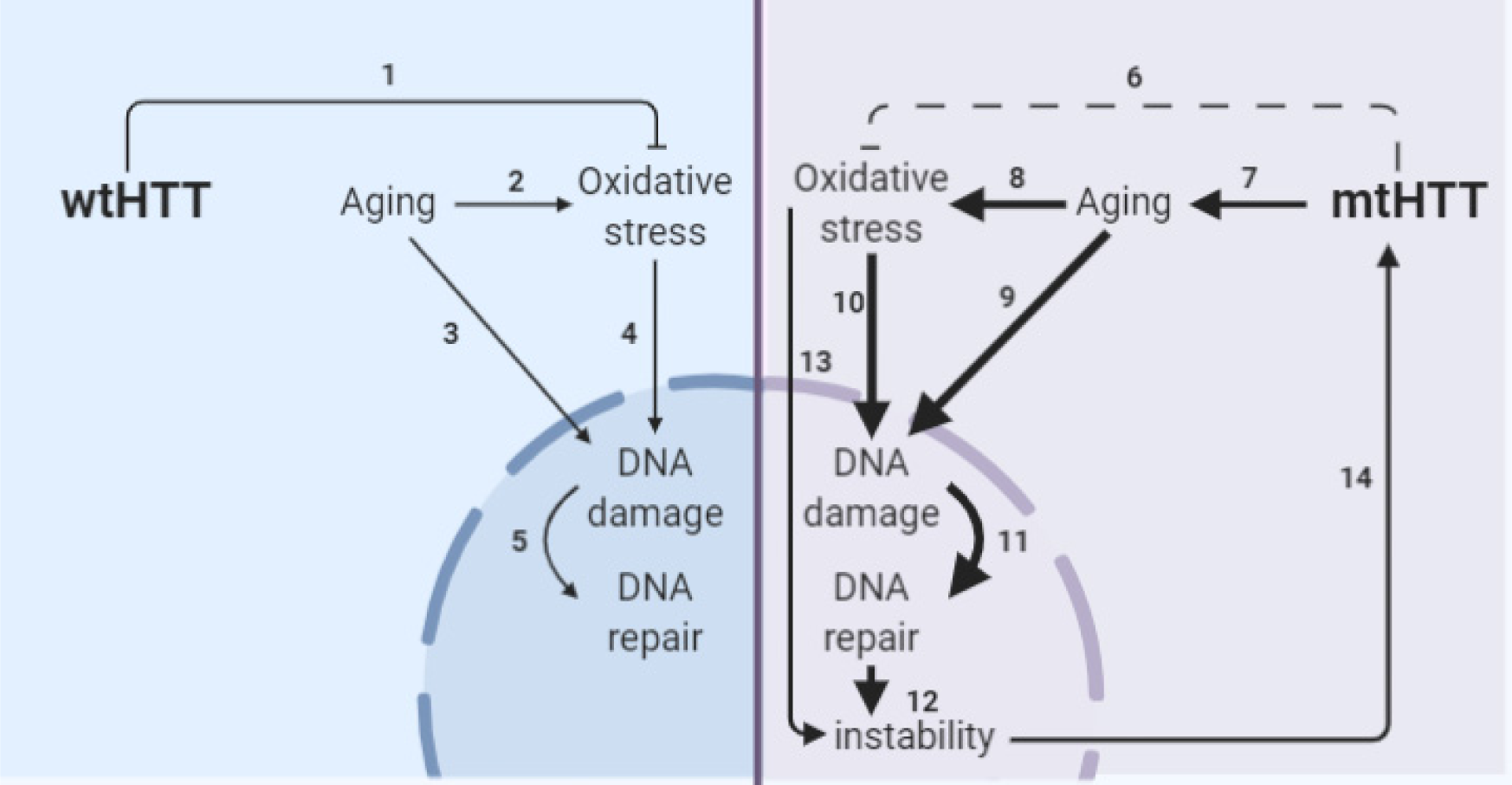
Model of the interplay of aging and oxidative stress in HD pathogenesis. (**1**) wtHTT is a stress response protein that can form Huntingtin stress bodies in the cytoplasm (Nath et al., 2015), can act as a ROS sensor (DiGiovanni, Mocle, Xia, & Truant, 2016), and aid in the base excision repair pathway in the nucleus in response to DNA damage (Maiuri et al., 2017). (**2**) Aging increases oxidative stress in cells (Andriollo-Sanchez et al., 2005; Beckman & Ames, 1998). (**3**) Aging increases DNA damage (Maynard, Fang, Scheibye-Knudsen, Croteau, & Bohr, 2015). (**4**) Oxidative stress increases oxidative damage, (Barzilai & Yamamoto, 2004). (**5**) DNA damage increases DNA repair particularly the base-excision repair pathway (Cadet, Davies, Medeiros, Di Mascio, & Wagner, 2017; Maynard et al., 2015; Prasad, Dyrkheeva, Williams, & Wilson, 2015). (**6**) mtHTT does not perform its stress response functions in the cell as well (Maiuri et al., 2017; Nath et al., 2015). Additionally, (**7**) accelerated epigenetic aging has been found in age-matched HD brains (Horvath et al., 2016) and age-related nuclear pore complex deficiencies have been found in HD (Grima et al., 2017), which collectively drive greater age-related oxidative stress and DNA damage and greater subsequent base excision DNA repair. (**8**) Altering the rate of base-excision repair can increase somatic instability of the CAG tract (Budworth et al., 2015; Goula et al., 2009; Kovtun et al., 2007). (**9**) Oxidative stress can cause somatic instability of the CAG tract (Jonson, Ougland, Klungland, & Larsen, 2013), possibly through the DNA damage/DNA repair pathway. (**10**) Somatic instability of the CAG tract is a driver of mtHTT toxicity, and may be necessary in certain tissues for onset and progression of HD (Budworth et al., 2015; Flower et al., 2019; Swami et al., 2009).

It should be noted that the DNA damage increase found in HD models has not been conclusively linked to aging (Goula et al., 2009), thus, further studies on this relationship are warranted with models of HD that possess an aging component. Additionally, future work should investigate directly if aging can modulate somatic instability and expansion in HD models. These findings have implications for potential modifiers of HD onset and/or progression within the aging process and suggest that anti-aging therapies and modulation of aging processes may be beneficial in treatment or prevention of HD.

## Materials and methods

### Primary Neuron Culture

Primary neurons were cultured from embryonic day (E)15.5-E17.5 humanized transgenic mice (Southwell et al., 2013). Brains were removed from embryos and stored in Hibernate-E (Thermo Fisher) overnight during genotyping. The following day, cortices with attached striatal tissue were dissected and pooled by genotype (Hu18/18 or Hu97/18). Cultures were established as in (Schmidt et al., 2018). Briefly, tissue was dissociated, typsinized, and resuspended in Neurobasal complete medium (NBC) (0.02% 10X SM1, 0.01% 100X Pen/Strep, 0.0025% glutamax in NeuroBasal Media (ThermoFisher Gibco 21103-049)). The cells were seeded onto poly-D-lysine (PDL) coated plates at a density of 250,000 cells per well in 24 well plates and 1×10^6^ in 6 well plates. For immunocytochemistry, coverslips were treated with hydrochloric acid overnight at room temperature (RT) with gentle shaking and washed in ethanol and phosphate buffered saline (PBS). Coverslips (Marinefield No. 1.5, 13mm) were then added to wells and dried prior to PDL coating. Cells were maintained in a 5% CO_2_/humidified incubator at 37°C. Neurons were fed with 1/10 well volume NBC twice weekly.

### Cell Treatment

For H_2_0_2_, staurosporine, and menadione treatments, a working solution of each oxidative stressor was added at 10x desired final concentration to NBC and given through a neuronal feed of 1/10 well volume. 30% H_2_O_2_ certified ACS stock (Fisher Chemical, H325-100) was diluted into pre-warmed NBC, and neurons were treated for 24h. 5 μM of a 72.4008 mM Menadione stock solution was made in PBS and added to pre-warmed NBC, and neurons were treated for 4h. A 1mM stock of staurosporine in DMSO was made, and neurons were treated with .5 μM in NBC for 1 hour. All conditions were performed in triplicate wells.

### Progerin Treatment

GFP-progerin cDNA was PCR amplified from the pBABE-puro-GFP-progerin plasmid (Addgene, 17663 deposited by Tom Misteli) using primers (forward 5’-GATCATCGATATGGTGAGCAAGGGCGAGGAG-3’, reverse 5’-GATCGCTAGCTTACATGATGCTGCAGTTCTGGG-3’). nGFP was PCR amplified from pAd nGFP (Addgene, 19413, deposited by Douglas Melton) using primers (forward 5’-CATCATCGAT ATGGTGCACGTGGATCCAA −3’, reverse 5’-GATCGCTAGCTTACTTGTACAGCTCGTCCA −3’). Amplified inserts were cloned into the ClaI and NheI sites of the AAV2 backbone vector (pFBAAVCAGmcsBgHpa, G0345) provided by the University of Iowa Viral Vector Core (UIVVC), which generated AAV2/1-GFP-progerin and AAV2/1-nGFP by the baculovirus system. Viral titers of 1.7e13 and 6e12 VG/ml for nGFP and progerin, respectively were obtained.

### Cell death

Cell death was measured in by release of lactate dehydrogenase (LDH) into the media and normalized to total remaining LDH in the lysate using the Pierce LDH Cytotoxicity Kit (Thermo Fisher, 88953) according to the manufacturer’s specifications.

### Reactive oxygen species detection

ROS was quantified using CM-H2DCFDA (Thermo Fisher Scientific catalog no. C6827) according to the manufacturer’s specifications. Briefly, CM-H2DCFDA in DMSO was diluted in NBC and added at a 5 μM concentration to each well. Neurons were protected from light and returned to the incubator for 30 minutes. In the last 5 minutes of incubation, Hoescht 3342 (Thermo Fisher, 62249) was added at 1/1000 well volume. After washing, ROS florescence was visualized and imaged with a Zeiss Axioplan fluorescence microscope. A minimum of 3 images per well were taken by a researcher blind to genotype and treatment, and integrated optical density (IOD) of fluorescence in images was quantified using ImageJ.

### Primary neuronal culture Western blot

Cells were lysed in the culture dishes in 1/10 well volume SDP lysis buffer (50 mM Tris pH 8.0, 150 mM Nacl, 1% Igepal/NP40, 40 mM B-gp (solid), 10 mMNaF and protease inhibitors: 50x Roche Complete, 1x NaVan, 1x PMSF, 1X zVAD). Dishes were incubated for 30 min with shaking and occasional hand agitation. Lysates were aspirated and cleared by centrifugation at 14,000 rpm for 10 minutes at 4°C and stored at −80°C until use. A DC assay (BioRad 5000111) was used to quantify lysate protein concentration. 35 ug of total protein was separated by SDS-PAGE using a 4-12% Bis Tris gel with MOPS running buffer (NuPage). Proteins were then transferred to 0.45um nitrocellulose membranes. Membranes were blocked in 5% dry milk power in PBS for 1 hour at RT and incubated overnight at 4°C in primary antibody (anti-Lamin B1 1µg/ml, Abcam ab16048, Cambridge, UK; anti-PCA NCAM 1:500, Millipore MAB5324, Burlington, MA; anti-Dynamin1 1 µg/ml, Abcam ab108458; anti-β-Tubulin 1:5000, Abcam ab131205; anti-β-Tubulin 1:5000, Abcam 6046) in 5% BSA in PBS-T. Proteins were detected with IR dye 800CW goat anti-mouse (1:250, Rockland 610-131-007, Gilbertsville, PA) and AlexaFluor 680 goat anti-rabbit (1:250, Molecular Probes A21076, Eugene, OR)-labeled secondary antibodies and the LiCor Odyssey Infrared Imaging system. Densitometry of bands was performed using ImageStudio 4.0 software (Li-Cor Biosciences). Lanes with abnormal loading control were excluded from analysis.

### YAC128 striatum Western blot

Striata from treated YAC128 animals were hand homogenized in lysis buffer containing 1% NP-40 detergent, a protease inhibitor cocktail (Roche, 4693159001), and a phosphatase inhibitor cocktail (Roche, 4906845001). 40µg of total protein was resolved on a 12% gel and transferred onto 0.45um nitrocellulose membrane. Blots were then blocked with 5% skim milk in PBS-T for 1 hour and incubated with either rabbit anti-lamin B1 (1:1000, Abcam, ab16048), rabbit anti-dynamin 1 (1:1000, Abcam, ab108458) or mouse anti-CDKN2A(p16INK4a) (1:1000, Abcam, AB54210) overnight at 4°C. Primary antibodies were detected with IR dye 800CW goat anti-mouse (1:5000, Rockland, 610-131-007) and AlexaFluor 680 goat anti-rabbit (1:5000, Molecular Probes, A21076)-labeled secondary antibodies. Blots were subsequently probed with mouse anti-β-Tubulin (1:5000 Abcam, ab6046) as a loading control and detected with IR dye 800CW goat anti-mouse. Proteins were visualized and quantified as above.

### iPSC differentiation and treatment

iPSC generation was previously performed at Cedar’s Sinai Medical Center using non-integrating methods from human fibroblast lines obtained from 3 HD patients with CAG repeat size of 43 (CS13iHD43n2), 71 (CS81iHD71n), or 109 (CS09iHD109n), and from 3 non-HD control subjects with CAG repeat sizes of 18 (CS25iCTR18n), 21 (CS00iCTR21n) or 33 (CS83iCTR33n). (HD iPSC Consortium, 2012; Mattis et al., 2015; Mehta et al., 2018). These iPSCs were fully reprogrammed as previously demonstrated by testing for pluripotency markers and gene expression analysis (Virginia B. Mattis et al., 2015), and removal of reprogramming plasmids was confirmed by genomic polymerase chain reaction (PCR) and southern blotting (Virginia B. Mattis et al., 2015). Neural stem cells (NSCs) were created using previously published protocols (Garcia et al., 2019; Telezhkin et al., 2016). NSCs were cryopreserved at 16 days of differentiation in DMSO-supplemented cell freezing media (Sigma, C6164) as described (Ebert et al., 2013). Coverslips were prepared by washing in HCl overnight, followed by washing 3x in ethanol and 3x in 1xPBS. Once dry, coverslips were transferred to the bottom of 24 well plates and coated with Corning® Matrigel® Matrix. NSCs were thawed onto coverslips, and directed toward a striatal fate by plating with neuronal induction media supplemented with GABA and pro-synaptogenic small molecules (CHIR99021 and forskolin), as previously published (Garcia et al., 2019; Kemp et al., 2016; Telezhkin et al., 2016). NSCs were infected with AAV2/1-nGFP or AAV2/1-progerin-GFP at 1e10 VG/ml the following day. Triplicate wells were used for each condition. Neurons were directed toward a striatal fate for an additional 9d with expression of progerin or nGFP, upon which neurons were fixed, stained, and transferred to slides using ProLong™ Gold Antifade Mountant with DAPI (Thermo Fisher). Two sets of neuron staining combinations were performed. The first set was stained for a marker of neuronal dendrites (MAP2 1:250, Sigma Aldrich m1408, St Louis, MO), a striatal neuron marker (DARPP-32 1:400, Cell Signaling 19A3, Beverly, MA), and viral transduction (GFP 1:2000, Abcam ab13970, Cambridge, UK). The second set was stained for a marker of apoptosis (cleaved caspase-3), DNA damage (γ-H_2_ AX 1:1000, Upstate Biotechnology), and viral transduction (GFP 1:2000, Abcam ab13970). Neurons were imaged using a Zeiss Axio Observer.Z1 confocal microscope at 40x and 63x objective magnification and were background-corrected using ImageJ with rolling ball radius of 35 pixels. Control (CAG18, CAG21, and CAG33) and HD (CAG43, CAG71, CAG109) were pooled for analyses.

### Dendritic analysis

Dendritic analysis was performed as described (Schmidt et al., 2018). Briefly, fluorescence images of cells stained with a marker of neuronal dendrites (MAP2 1:250, Sigma Aldrich m1408) were acquired using a Leica TCS SP8 confocal laser scanning microscope at 63X objective magnification. Samples from different groups were interleaved and the researcher was blinded to experimental conditions during imaging and analysis. Image stacks of Z-step size of 60 μm were converted to 2D in Image J using the maximum intensity Z-projection function. Images were then background subtracted with a rolling ball radius of 35 pixels and de-speckled. Images were imported into NeuronStudio (Version 0.9.92) for semi-automated measurement of dendritic length and Sholl analysis.

### Immunocytochemistry

Neurons grown on coverslips were fixed with 4% PFA for 20min at room temp, diluted in 1xPBS, washed in 1xPBS, and blocked in 3% BSA, 5% normal goat serum and .15% triton in PBS for 20 minutes at RT. Fixed cells were then incubated overnight at 4°C in primary antibody (GFP 1:2000, Abcam ab13970, γH2A.X 1:500, Sigma Aldrich 05-636; Map-2 1:250, Sigma Aldrich M4403; Map-2 1:250, Invitrogen PA517646; EM48 1:100, Millipore MAB5374; LC3B 1:2000, Novus Biologicals NB600-1384 Centennial, CO; DNA/RNA damage [15A3] 1:500 Abcam ab62623; DARPP-32 1:500, Abcam ab40801; Cleaved caspase-3 1:400, Cell Signal 9664 Danvers, MA; TUJ-1 1:1000, Sigma Aldrich T8660; Nestin 1:500 Millipore MAB5326), washed 3x with PBS, and subsequently incubated in secondary antibody (goat anti-chicken 488 1:200, Invitrogen A32931; goat anti-rabbit 647 1:200, Invitrogen A21244; goat anti-rabbit 568 1:200, Invitrogen A11011; goat anti-mouse 647, Invitrogen A21235 1:200; goat anti-mouse 568 1:200, Invitrogen A11004) for 1 hour at RT protected from light. After washing 3x with PBS, coverslips were mounted on 75mm slides (Fisherbrand, 12-550-15) with ProLong Gold Antifade mounting reagent with DAPI. Stained Cells were imaged with a Zeiss Axioplan fluorescence microscope with 20x, 40x, and 63x magnification objectives. At least 3 images were taken per well and integrated density/fluorescence was quantified using ImageJ.

### Surgical AAV delivery

AAVs were delivered by stereotaxic intrastriatal convection enhanced delivery as in (Southwell et al., 2009). Briefly, a burr hole was made at 0.8 mm anterior and 2 mm lateral to Bregma. A Hamilton syringe with 30-gauge needle preloaded with 4µl sterile saline and 1 µl virus was slowly lowered to 3.5 mm below the dura. A microinjector was then used to inject the virus followed by the bolus of sterile saline at a rate of 0.5 µl/min. The needs was left in place for 5 minutes and slowly withdrawn.

### Tissue collection and processing

For mice allotted for terminal molecular and biochemical analysis, brains were removed and placed on ice for 1 min to increase tissue rigidity. Brains were then microdissected by region. Striata and cortices were preserved in RNAlater (Ambion) overnight at 4°C and then stored at - 80°C until use. Mice allotted for terminal histological analysis were perfused transcardially with PBS and 4% PFA. Brains were removed and post-fixed in 4% PFA in PBS for 24 hrs at 4°C. The following day brains were cryoprotected in 30% sucrose with 0.01% sodium azide. Once equilibrated, brains were divided into forebrain and cerebellum, and forebrains were frozen on dry ice, mounted in Tissue-TEK O.C.T. embedding compound (Sakura), and cut via cryostat (Leica CM3050S) into a series of 25 μm coronal sections free-floating in PBS with 0.01% sodium azide.

### Brain histology

Sections were stained with primary rabbit anti-GFP (1:1000, Life Technologies) mouse anti-DNA/RNA damage antibody [15A3] (1:500, Abcam ab62623) and secondary goat anti-rabbit (1:500, Alexa-fluor 488) and goat-anti-mouse (1:500, Alexa Fluor 568). Sections were mounted using ProLong Gold Antifade mounting reagent with DAPI. Sections were imaged with a 2.5x objective (ZEISS Microscopy) using a Zeiss Axio Vert.A1 microscope (ZEISS Microscopy), Zeiss AxioCam ICm1 camera (Carl Zeiss) and Zeiss Zen 2.3 Lite software (Carl Zeiss). 4 images per section were merged using the PhotoMerge tool in Adobe Photoshop CC 2018 (Adobe).

### Statistical Analysis

Statistical analyses were performed in Graphpad Prism (v. 8.1.1). 2 way ANOVA with Bonferroni *post-hoc* correction was performed to determine differences between treatments and genotypes or unpaired t-tests for comparison of only 2 groups. For image quantification, images with fewer than 3 neurons (determined by DAPI quantification) were excluded from analyses.

## Acknowledgements

The authors would like to thank Jakob Rupar, Casey Hart, Katlin Hencak, Mark Wang, and Qingwen Xia, for assistance with mouse husbandry and genotyping, Christopher Yanick and Christopher Parrett for assistance with cloning, Addgene for providing pBABE-puro-GFP-progerin (17663) and pAd-nGFP (19413) plasmids, and the University of Iowa Viral Vector Core for AAV production.

## References

Andrew, S. E., Goldberg, Y. P., Kremer, B., Telenius, H., Theilmann, J., Adam, S., Starr, E., Squitieri, F., Lin, B., Kalchman, M., Graham, R., & Hayden, M. R. (1993). The relationship between trinucleotide (CAG) repeat length and clinical features of Huntington’s disease. Nature Genetics, 4(4), 398–403. https://doi.org/10.1038/ng0893-398

Andriollo-Sanchez, M., Hininger-Favier, I., Meunier, N., Venneria, E., O’Connor, J. M., Maiani, G., Coudray, C., & Roussel, A. M. (2005). Age-related oxidative stress and antioxidant parameters in middle-aged and older European subjects: the ZENITH study. European Journal of Clinical Nutrition, 59(2): S58–62. doi:10.1038/sj.ejcn.1602300

Atwal, R. S., Xia, J., Pinchev, D., Taylor, J., Epand, R. M., & Truant, R. (2007). Huntingtin has a membrane association signal that can modulate huntingtin aggregation, nuclear entry and toxicity. Hum Mol Genet, 16(21), 2600–2615. doi:10.1093/hmg/ddm217

Baker, D. J., Jin, F., & van Deursen, J. M. (2008). The yin and yang of the Cdkn2a locus in senescence and aging. Cell Cycle (Georgetown, Tex.), 7(18), 2795–2802.

Bates, G. P., Dorsey, R., Gusella, J. F., Hayden, M. R., Kay, C., Leavitt, B. R., … Tabrizi, S. J. (2015). Huntington disease. Nat Rev Dis Primers, 1, 15005. doi:10.1038/nrdp.2015.5

Beckhauser, T. F., Francis-Oliveira, J., & De Pasquale, R. (2016). Reactive oxygen species: physiological and physiopathological effects on synaptic plasticity. Journal of Experimental Neuroscience, 10(Suppl 1), 23–48. https://doi.org/10.4137/JEN.S39887

Beckman, K. B., & Ames, B. N. (1998). The free radical theory of aging matures. Physiological Review, 78(2), 547–581. doi:10.1152/physrev.1998.78.2.547

Biffi, E., Regalia, G., Menegon, A., Ferrigno, G., & Pedrocchi, A. (2013). The influence of neuronal density and maturation on network activity of hippocampal cell cultures: A methodological study. PLoS ONE, 8(12). https://doi.org/10.1371/journal.pone.0083899

Bogdanov, M. B., Andreassen, O. A., Dedeoglu, A., Ferrante, R. J., & Beal, M. F. (2001). Increased oxidative damage to DNA in a transgenic mouse model of Huntington’s disease. Journal of Neurochemistry, 79(6), 1246–1249. doi:10.1046/j.1471-4159.2001.00689.x

Barzilai, A., & Yamamoto, K. (2004). DNA damage responses to oxidative stress. DNA Repair (Amst), 3(8-9), 1109–1115. doi:10.1016/j.dnarep.2004.03.002

Brocardo, P. S., McGinnis, E., Christie, B. R., & Gil-Mohapel, J. (2016). Time-course analysis of protein and lipid oxidation in the brains of YAC128 Huntington’s disease transgenic mice. Rejuvenation Research, 19(2), 140–148. https://doi.org/10.1089/rej.2015.1736

Browne, S. E., & Beal, M. F. (2006). Oxidative damage in Huntington’s disease pathogenesis. Antioxid Redox Signal, 8(11-12), 2061–2073. doi:10.1089/ars.2006.8.2061

Browne, S. E., Ferrante, R. J., & Beal, M. F. (1999). Oxidative stress in Huntington’s disease. Brain Pathol, 9(1), 147–163.

Budworth, H., Harris, F. R., Williams, P., Lee, D. Y., Holt, A., Pahnke, J., … McMurray, C. T. (2015). Suppression of Somatic Expansion Delays the Onset of Pathophysiology in a Mouse Model of Huntington’s Disease. PLoS Genetics, 11(8): e1005267. doi:10.1371/journal.pgen.1005267

Buren, C., Tu, G., Parsons, M. P., Sepers, M. D., & Raymond, L. A. (2016). Influence of cortical synaptic input on striatal neuronal dendritic arborization and sensitivity to excitotoxicity in corticostriatal coculture. Journal of Neurophysiology, 116(2), 380–390. https://doi.org/10.1152/jn.00933.2015

Cadet, J., Davies, K. J. A., Medeiros, M. H., Di Mascio, P., & Wagner, J. R. (2017). Formation and repair of oxidatively generated damage in cellular DNA. Free Radic Biol Med, 107, 13–34. doi:10.1016/j.freeradbiomed.2016.12.049

Cao, K., Blair, C. D., Faddah, D. A., Kieckhaefer, J. E., Olive, M., Erdos, M. R., Nabel, EG., & Collins, F. S. (2011). Progerin and telomere dysfunction collaborate to trigger cellular senescence in normal human fibroblasts. The Journal of Clinical Investigation, 121(7), 2833–2844. https://doi.org/10.1172/JCI43578

Carroll, J. B., Lerch, J. P., Franciosi, S., Spreeuw, A., Bissada, N., Henkelman, R. M., & Hayden, M. R. (2011). Natural history of disease in the YAC128 mouse reveals a discrete signature of pathology in Huntington disease. Neurobiology of disease, 43(1), 257–265.

Chen, C. M., Wu, Y. R., Cheng, M. L., Liu, J. L., Lee, Y. M., Lee, P. W., … Chiu, D. T. (2007). Increased oxidative damage and mitochondrial abnormalities in the peripheral blood of Huntington’s disease patients. Biochem Biophys Res Commun, 359(2), 335–340. doi:10.1016/j.bbrc.2007.05.093

Chen, L., Liu, L., Yin, J., Luo, Y., & Huang, S. (2009). Hydrogen peroxide-induced neuronal apoptosis is associated with inhibition of protein phosphatase 2A and 5, leading to activation of MAPK pathway. The International Journal of Biochemistry & Cell Biology, 41(6), 1284–1295. https://doi.org/10.1016/j.biocel.2008.10.029

Consortium, H. D. i. (2012). Induced pluripotent stem cells from patients with Huntington’s disease show CAG-repeat-expansion-associated phenotypes. Cell Stem Cell, 11(2), 264–278. doi:10.1016/j.stem.2012.04.027

Croce, K. R., & Yamamoto, A. (2019). A role for autophagy in Huntington’s disease. Neurobiology of Disease, 122, 16–22. https://doi.org/10.1016/j.nbd.2018.08.010

Dickey, A. S., Sanchez, D. N., Arreola, M., Sampat, K. R., Fan, W., Arbez, N., Akimov, S., Van Kanegan, MJ., Ohnishi, K., Gilmore-Hall, SK., Flores, AL., Nguyen, JM., Lomas, N, Hsu, CL., Lo, DC., Ross, CA., Masliah, E., Evans, RM., & Spada, A. R. L. (2017). PPARd activation by bexarotene promotes neuroprotection by restoring bioenergetic and quality control homeostasis. Science Translational Medicine, 9(419), eaal2332. https://doi.org/10.1126/scitranslmed.aal2332

DiGiovanni, L. F., Mocle, A. J., Xia, J., & Truant, R. (2016). Huntingtin N17 domain is a reactive oxygen species sensor regulating huntingtin phosphorylation and localization. Hum Mol Genet, 25(18), 3937–3945. doi:10.1093/hmg/ddw234

Diguet, E., Petit, F., Escartin, C., Cambon, K., Bizat, N., Dufour, N., Hantraye, P., Deglon, N., & Brouillet, E. (2009). Normal aging modulates the neurotoxicity of mutant Huntingtin. PLOS ONE, 4(2), e4637. https://doi.org/10.1371/journal.pone.0004637

Dragileva, E., Hendricks, A., Teed, A., Gillis, T., Lopez, E. T., Friedberg, E. C., … Wheeler, V. C. (2009). Intergenerational and striatal CAG repeat instability in Huntington’s disease knock-in mice involve different DNA repair genes. Neurobiol Dis, 33(1), 37–47. doi:10.1016/j.nbd.2008.09.014

Dumas, E. M., van den Bogaard, S. J. A., Ruber, M. E., Reilman, R. R., Stout, J. C., Craufurd, D., Hicks, SL., Kennard, C., Tabrizi, SJ., van Buchem, MA., van der Grond, J., & Roos, R. A. C. (2012). Early changes in white matter pathways of the sensorimotor cortex in premanifest Huntington’s disease. Human Brain Mapping, 33(1), 203–212. https://doi.org/10.1002/hbm.21205

Ebert, A. D., Shelley, B. C., Hurley, A. M., Onorati, M., Castiglioni, V., Patitucci, T. N., … Svendsen, C. N. (2013). EZ spheres: A stable and expandable culture system for the generation of pre-rosette multipotent stem cells from human ESCs and iPSCs. Stem Cell Research, 10(3), 417–427.

Ehrnhoefer, D. E., Martin, D. D. O., Schmidt, M. E., Qiu, X., Ladha, S., Caron, N. S., Skotte, NH., Nguyen, YTN., Vaid, K., Southwell, AL., Engemann, S., Franciosi, S., & Hayden, MR. (2018). Preventing mutant huntingtin proteolysis and intermittent fasting promote autophagy in models of Huntington disease. Acta Neuropathologica Communications, 6(1), 16. https://doi.org/10.1186/s40478-018-0518-0

Eriksson, M., Brown, W. T., Gordon, L. B., Glynn, M. W., Singer, J., Scott, L., Erdos, MR., Robbins, CM., Moses, TY., Berglund, P., Dutra, A., Pak, E., Durkin, S., Csoka, AB., Boehnke, M., Glover, TW., & Collins, F. S. (2003). Recurrent de novo point mutations in lamin A cause Hutchinson–Gilford progeria syndrome. Nature, 423(6937), 293–298. https://doi.org/10.1038/nature01629

Flower, M., Lomeikaite, V., Ciosi, M., Cumming, S., Morales, F., Lo, K., … Tabrizi, S. J. (2019). MSH3 modifies somatic instability and disease severity in Huntington’s and myotonic dystrophy type 1. Brain. doi:10.1093/brain/awz115

Freund, A., Laberge, R.-M., Demaria, M., & Campisi, J. (2012). Lamin B1 loss is a senescence-associated biomarker. Molecular Biology of the Cell, 23(11), 2066–2075. https://doi.org/10.1091/mbc.E11-10-0884

Garcia, V. J., Rushton, D. J., Tom, C. M., Allen, N. D., Kemp, P. J., Svendsen, C. N., & Mattis, V. B. (2019). Huntington’s disease patient-derived astrocytes display electrophysiological impairments and reduced neuronal support. Frontiers in Neuroscience, 13. https://doi.org/10.3389/fnins.2019.00669

Gazzaley, A. H., Weiland, N. G., McEwen, B. S., & Morrison, J. H. (1996). Differential regulation of NMDAR1 mRNA and protein by estradiol in the rat hippocampus. Journal of Neuroscience, 16(21), 6830–6838. https://doi.org/10.1523/JNEUROSCI.16-21-06830.1996

Genetic Modifiers of Huntington’s Disease, C. (2015). Identification of Genetic Factors that Modify Clinical Onset of Huntington’s Disease. Cell, 162(3), 516–526. doi:10.1016/j.cell.2015.07.003

Gorbunova, V., Seluanov, A., Mao, Z., & Hine, C. (2007). Changes in DNA repair during aging. Nucleic Acids Research, 35(22), 7466–7474. https://doi.org/10.1093/nar/gkm756

Goula, A. V., Berquist, B. R., Wilson, D. M., 3rd, Wheeler, V. C., Trottier, Y., & Merienne, K. (2009). Stoichiometry of base excision repair proteins correlates with increased somatic CAG instability in striatum over cerebellum in Huntington’s disease transgenic mice. PLoS Genet, 5(12), e1000749. doi:10.1371/journal.pgen.1000749

Graham, R. K., Pouladi, M. A., Joshi, P., Lu, G., Deng, Y., Wu, N.-P., Figueroa, BE., Metzler, M., Andre, VM., Slow, EJ., Raymond, L., Friedlander, R., Levine, MS., Leavitt, BR., & Hayden, MR. (2009). Differential susceptibility to excitotoxic stress in YAC128 mouse models of Huntington disease between initiation and progression of disease. Journal of Neuroscience, 29(7), 2193–2204. https://doi.org/10.1523/JNEUROSCI.5473-08.2009

Graveland, G. A., Williams, R. S., & DiFiglia, M. (1985). Evidence for degenerative and regenerative changes in neostriatal spiny neurons in Huntington’s disease. Science (New York, N.Y.), 227(4688), 770–773. https://doi.org/10.1126/science.3155875

Grima, J. C., Daigle, J. G., Arbez, N., Cunningham, K. C., Zhang, K., Ochaba, J., Geater, C., Morozko, E., Stocksdale, J., Glatzer, JC., Pham, JT., Ahmed, I., Peng, Q., Wadhwa, H., Pletnikova, O., Troncoso, JC., Duan, W., Snyder, SH., Ranum, LPW., Thompson, LM., Lloyd, TE., Ross, CA., & Rothstein, JD. (2017). Mutant Huntingtin disrupts the nuclear pore complex. Neuron, 94(1), 93–107.e6. https://doi.org/10.1016/j.neuron.2017.03.023

Gump, J. M., & Thorburn, A. (2011). Autophagy and apoptosis-what’s the connection? Trends in Cell Biology, 21(7), 387–392. https://doi.org/10.1016/j.tcb.2011.03.007

HD Coll Res Grp. (1993). A novel gene containing a trinucleotide repeat that is expanded and unstable on Huntington’s disease chromosomes. The Huntington’s Disease Collaborative Research Group. Cell, 72(6), 971–983. doi:10.1016/0092-8674(93)90585-e

HD iPSC Consortium. (2012). Induced pluripotent stem cells from patients with Huntington’s disease show CAG-repeat-expansion-associated phenotypes. Cell Stem Cell, 11(2), 264–278. https://doi.org/10.1016/j.stem.2012.04.027

Hermel, E., Gafni, J., Propp, S. S., Leavitt, B. R., Wellington, C. L., Young, J. E., Hackam, AS., Logvinova, AV., Peel, AL., Chen, SF., Hook, V., Singaraja, R., Krajewski, S., Goldsmith, PC., Ellerby, HM., Hayden, MR., Bredesen, DE., & Ellerby, LM. (2004). Specific caspase interactions and amplification are involved in selective neuronal vulnerability in Huntington’s disease. Cell Death & Differentiation, 11(4), 424–438. https://doi.org/10.1038/sj.cdd.4401358

Higgins, G. C., Devenish, R. J., Beart, P. M., & Nagley, P. (2011). Autophagic activity in cortical neurons under acute oxidative stress directly contributes to cell death. Cellular and Molecular Life Sciences, 68(22), 3725–3740. https://doi.org/10.1007/s00018-011-0667-9

Hodgson, J. G., Agopyan, N., Gutekunst, C. A., Leavitt, B. R., LePiane, F., Singaraja, R., Smith, DJ., Bissada, N., McCutcheon, K., Nasir, J., Jamot, L., Li, XJ., Stevens, ME., Rosemond, E., Order, JC., Phillips, AG., Rubin, EM., Hersch, SM., & Hayden, MR. (1999). A YAC mouse model for Huntington’s disease with full-length mutant huntingtin, cytoplasmic toxicity, and selective striatal neurodegeneration. Neuron, 23(1), 181–192. https://doi.org/10.1016/s0896-6273(00)80764-3

Horvath, S., Langfelder, P., Kwak, S., Aaronson, J., Rosinski, J., Vogt, T. F., Eszes, M., Faull, R., Curtis, M., Waldvogel., H., Choi, O., Tung, S., Vinters, H., Coppola, G., & Yang, XW. (2016). Huntington’s disease accelerates epigenetic aging of human brain and disrupts DNA methylation levels. Aging (Albany NY), 8(7), 1485–1504. https://doi.org/10.18632/aging.101005

Jin, Y. N., Hwang, W. Y., Jo, C., & Johnson, G. V. W. (2012). Metabolic state determines sensitivity to cellular stress in Huntington disease: normalization by activation of PPARγ. PLOS ONE, 7(1), e30406. https://doi.org/10.1371/journal.pone.0030406

Johri, A., & Beal, M. F. (2012). Antioxidants in Huntington’s disease. Biochimica Et Biophysica Acta, 1822(5), 664–674. https://doi.org/10.1016/j.bbadis.2011.11.014

Jonson, I., Ougland, R., Klungland, A., & Larsen, E. (2013). Oxidative stress causes DNA triplet expansion in Huntington’s disease mouse embryonic stem cells. Stem Cell Research, 11(3), 1264–1271. https://doi.org/10.1016/j.scr.2013.08.010

Kemp, P. J., Rushton, D. J., Yarova, P. L., Schnell, C., Geater, C., Hancock, J. M., Wieland, A., Hughes, A., Badder, L., Cope, E., Riccardi, D., Randall, AD., Brown, JT., Allen, ND., & Telezhkin, V. (2016). Improving and accelerating the differentiation and functional maturation of human stem cell-derived neurons: Role of extracellular calcium and GABA. The Journal of Physiology, 594(22), 6583–6594. https://doi.org/10.1113/JP270655

Kim, J., Moody, J. P., Edgerly, C. K., Bordiuk, O. L., Cormier, K., Smith, K., Beal, MF., & Ferrante, RJ. (2010). Mitochondrial loss, dysfunction and altered dynamics in Huntington’s disease. Human Molecular Genetics, 19(20), 3919–3935. https://doi.org/10.1093/hmg/ddq306

Klepac, N., Relja, M., Klepac, R., Hecimovic, S., Babic, T., & Trkulja, V. (2007). Oxidative stress parameters in plasma of Huntington’s disease patients, asymptomatic Huntington’s disease gene carriers and healthy subjects: a cross-sectional study. J Neurol, 254(12), 1676–1683. doi:10.1007/s00415-007-0611-y

Kovtun, I. V., Liu, Y., Bjoras, M., Klungland, A., Wilson, S. H., & McMurray, C. T. (2007). OGG1 initiates age-dependent CAG trinucleotide expansion in somatic cells. Nature, 447(7143), 447–452. https://doi.org/10.1038/nature05778

Kruman, I., Guo, Q., & Mattson, M. P. (1998). Calcium and reactive oxygen species mediate staurosporine-induced mitochondrial dysfunction and apoptosis in PC12 cells. Journal of Neuroscience Research, 51(3), 293–308. https://doi.org/10.1002/(SICI)1097-4547(19980201)51:3<293::AID-JNR3>3.0.CO;2-B

Lai, Y., Budworth, H., Beaver, J. M., Chan, N. L., Zhang, Z., McMurray, C. T., & Liu, Y. (2016). Crosstalk between MSH2-MSH3 and polbeta promotes trinucleotide repeat expansion during base excision repair. Nature Communications, 7, 12465. doi:10.1038/ncomms12465

Lee, S., Sato, Y., & Nixon, R. A. (2011). Lysosomal proteolysis inhibition selectively disrupts axonal transport of degradative organelles and causes an Alzheimer’s-like axonal dystrophy. The Journal of Neuroscience, 31(21), 7817–7830. https://doi.org/10.1523/JNEUROSCI.6412-10.2011

Lerner, R. P., Trejo, C. M. L., Zhu, C., Chesselet, M. F., & Hickey, M. A. (2012). Striatal atrophy and dendritic alterations in a knock-in mouse model of Huntington’s disease. Brain Research Bulletin, 87(6), 571–578. https://doi.org/10.1016/j.brainresbull.2012.01.012

Li, S.-H., Lam, S., Cheng, A. L., & Li, X.-J. (2000). Intranuclear huntingtin increases the expression of caspase-1 and induces apoptosis. Human Molecular Genetics, 9(19), 2859–2867. https://doi.org/10.1093/hmg/9.19.2859

Loor, G., Kondapalli, J., Schriewer, J. M., Chandel, N. S., Vanden Hoek, T. L., & Schumacker, P. T. (2010). Menadione triggers cell death through ROS-dependent mechanisms involving PARP activation without requiring apoptosis. Free Radical Biology & Medicine, 49(12), 1925–1936. https://doi.org/10.1016/j.freeradbiomed.2010.09.021

Maiuri, T., Mocle, A. J., Hung, C. L., Xia, J., van Roon-Mom, W. M. C., & Truant, R. (2017). Huntingtin is a scaffolding protein in the ATM oxidative DNA damage response complex. Human Molecular Genetics, 26(2), 395–406. https://doi.org/10.1093/hmg/ddw395

Manley, K., Shirley, T. L., Flaherty, L., & Messer, A. (1999). Msh2 deficiency prevents in vivo somatic instability of the CAG repeat in Huntington disease transgenic mice. Nature Genetics, 23(4), 471–473. https://doi.org/10.1038/70598

Martin, D. D. O., Ladha, S., Ehrnhoefer, D. E., & Hayden, M. R. (2015). Autophagy in Huntington disease and huntingtin in autophagy. Trends in Neurosciences, 38(1), 26–35. https://doi.org/10.1016/j.tins.2014.09.003

Martinez-Vicente, M., Talloczy, Z., Wong, E., Tang, G., Koga, H., Kaushik, S., … Cuervo, A. M. (2010). Cargo recognition failure is responsible for inefficient autophagy in Huntington’s disease. Nature Neuroscience, 13(5), 567–576. doi:10.1038/nn.2528

Mattis, V. B., Tom, C., Akimov, S., Saeedian, J., Oestergaard, M. E., Southwell, A. L., … Svendsen, C. N. (2015). HD iPSC-derived neural progenitors accumulate in culture and are susceptible to BDNF withdrawal due to glutamate toxicity. Human Molecular Genetics, 24(11), 3257–3271. doi:10.1093/hmg/ddv080

Maynard, S., Fang, E. F., Scheibye-Knudsen, M., Croteau, D. L., & Bohr, V. A. (2015). DNA Damage, DNA Repair, Aging, and Neurodegeneration. Cold Spring Harb Perspect Med, 5(10). doi:10.1101/cshperspect.a025130

McClintock, D., Ratner, D., Lokuge, M., Owens, D. M., Gordon, L. B., Collins, F. S., & Djabali, K. (2007). The mutant form of lamin A that causes Hutchinson-Gilford progeria is a biomarker of cellular aging in human skin. PloS One, 2(12), e1269. https://doi.org/10.1371/journal.pone.0001269

Mehta, S. R., Tom, C. M., Wang, Y., Bresee, C., Rushton, D., Mathkar, P. P., Tang, J., & Mattis, VB. (2018). Human Huntington’s disease iPSC-derived cortical neurons display altered transcriptomics, morphology, and maturation. Cell Reports, 25(4), 1081–1096.e6. https://doi.org/10.1016/j.celrep.2018.09.076

Miller, J. D., Ganat, Y. M., Kishinevsky, S., Bowman, R. L., Liu, B., Tu, E. Y., … Studer, L. (2013). Human iPSC-based modeling of late-onset disease via progerin-induced aging. Cell Stem Cell, 13(6), 691–705. doi:10.1016/j.stem.2013.11.006

Moss, D. J. H., Pardinas, A. F., Langbehn, D., Lo, K., Leavitt, B. R., Roos, R., … Tabrizi, S. J. (2017). Identification of genetic variants associated with Huntington’s disease progression: a genome-wide association study. Lancet Neurol, 16(9), 701–711. doi:10.1016/S1474-4422(17)30161-8

Nance, M. A., & Myers, R. H. (2001). Juvenile onset Huntington’s disease—Clinical and research perspectives. Mental Retardation and Developmental Disabilities Research Reviews, 7(3), 153–157. https://doi.org/10.1002/mrdd.1022

Nath, S., Munsie, L. N., & Truant, R. (2015). A huntingtin-mediated fast stress response halting endosomal trafficking is defective in Huntington’s disease. Human Molecular Genetics, 24(2), 450–462. https://doi.org/10.1093/hmg/ddu460

Peña-Sánchez, M., Riverón-Forment, G., Zaldívar-Vaillant, T., Soto-Lavastida, A., Borrero-Sánchez, J., Lara-Fernández, G., Esteban-Hernandez, EM., Hernandez-Diaz, Z., Gonzalez-Quevedo, A., Fernandez-Almirall, I., Perez-Lopez, C., Castillo-Casanas, Y., Martinez-Bonne, O., Cabrera-Rivero, A., Valdes-Ramos, L., Guerra-Badia, R., Fernandez-Carriera, R., Menendez-Sainz, MC., & González-García, S. (2015). Association of status redox with demographic, clinical and imaging parameters in patients with Huntington’s disease. Clinical Biochemistry, 48(18), 1258–1263. https://doi.org/10.1016/j.clinbiochem.2015.06.014

Pham-Huy, L. A., He, H., & Pham-Huy, C. (2008). Free radicals, antioxidants in disease and health. International Journal of Biomedical Science: IJBS, 4(2), 89–96.

Polidori, M. C., Mecocci, P., Browne, S. E., Senin, U., & Beal, M. F. (1999). Oxidative damage to mitochondrial DNA in Huntington’s disease parietal cortex. Neuroscience Letters, 272(1), 53–56. https://doi.org/10.1016/S0304-3940(99)00578-9

Polyzos, A., Holt, A., Brown, C., Cosme, C., Wipf, P., Gomez-Marin, A., Castro, MR., Ayala-Pena, S., & McMurray, C. T. (2016). Mitochondrial targeting of XJB-5-131 attenuates or improves pathophysiology in HdhQ150 animals with well-developed disease phenotypes. Human Molecular Genetics, 25(9), 1792–1802. https://doi.org/10.1093/hmg/ddw051

Prasad, R., Dyrkheeva, N., Williams, J., & Wilson, S. H. (2015). Mammalian Base Excision Repair: Functional Partnership between PARP-1 and APE1 in AP-Site Repair. Plos One, 10(5), e0124269. doi:10.1371/journal.pone.0124269

Rotblat, B., Southwell, A. L., Ehrnhoefer, D. E., Skotte, N. H., Metzler, M., Franciosi, S., Leprivier, G., Somasekharan, SP., Barokas, A., Deng, Y., Tang, T., Mathers, J., Cetinbas, N., Daugaard, M., Kwok, B., Li, L., Carnie, CJ., Fink, D., Nitsch, R., Galpin, JD., Ahern, CA., Melino, G., Penninger, JM., Hayden, MR., & Sorensen, P. H. (2014). HACE1 reduces oxidative stress and mutant Huntingtin toxicity by promoting the NRF2 response. Proceedings of the National Academy of Sciences, 111(8), 3032–3037. https://doi.org/10.1073/pnas.1314421111

Schieber, M., & Chandel, N. S. (2014). ROS function in redox signaling and oxidative stress. Current Biology, 24(10), R453–462. doi:10.1016/j.cub.2014.03.034

Schmidt, M. E., Buren, C., Mackay, J. P., Cheung, D., Dal Cengio, L., Raymond, L. A., & Hayden, M. R. (2018). Altering cortical input unmasks synaptic phenotypes in the YAC128 cortico-striatal coculture model of Huntington disease. BMC Biol, 16(1), 58. doi:10.1186/s12915-018-0526-3

Sepers, M. D., & Raymond, L. A. (2014). Mechanisms of synaptic dysfunction and excitotoxicity in Huntington’s disease. Drug Discovery Today, 19(7), 990–996. https://doi.org/10.1016/j.drudis.2014.02.006

Skotte, N. H., Southwell, A. L., Ostergaard, M. E., Carroll, J. B., Warby, S. C., Doty, C. N., … Hayden, M. R. (2014). Allele-specific suppression of mutant huntingtin using antisense oligonucleotides: Providing a therapeutic option for all Huntington disease patients. PLoS ONE, 9(9), e107434.

Slow, E. J., van Raamsdonk, J., Rogers, D., Coleman, S. H., Graham, R. K., Deng, Y., … Hayden, M. R. (2003). Selective striatal neuronal loss in a YAC128 mouse model of Huntington disease. Hum Mol Genet, 12(13), 1555–1567.

Snell, R. G., MacMillan, J. C., Cheadle, J. P., Fenton, I., Lazarou, L. P., Davies, P., MacDonald, ME., Gusella, JF., Harper, PS., & Shaw, D. J. (1993). Relationship between trinucleotide repeat expansion and phenotypic variation in Huntington’s disease. Nature Genetics, 4(4), 393–397. https://doi.org/10.1038/ng0893-393

Sorolla, M. A., Reverter-Branchat, G., Tamarit, J., Ferrer, I., Ros, J., & Cabiscol, E. (2008). Proteomic and oxidative stress analysis in human brain samples of Huntington disease. Free Radical Biology & Medicine, 45(5), 667–678. https://doi.org/10.1016/j.freeradbiomed.2008.05.014

Southwell, A. L., Ko, J., & Patterson, P. H. (2009). Intrabody gene therapy ameliorates motor, cognitive, and neuropathological symptoms in multiple mouse models of Huntington’s disease. J. Neurosci., 29(43), 13589–13602. doi:10.1523/jneurosci.4286-09.2009

Southwell, A. L., Warby, S. C., Carroll, J. B., Doty, C. N., Skotte, N. H., Zhang, W., … Hayden, M. R. (2013). A fully humanized transgenic mouse model of Huntington disease. Human Molecular Genetics, 22(1), 18–34. doi:10.1093/hmg/dds397

Swami, M., Hendricks, A. E., Gillis, T., Massood, T., Mysore, J., Myers, R. H., & Wheeler, V. C. (2009). Somatic expansion of the Huntington’s disease CAG repeat in the brain is associated with an earlier age of disease onset. Hum Mol Genet, 18(16), 3039–3047. doi:10.1093/hmg/ddp242

Tabrizi, S. J., Scahill, R. I., Owen, G., Durr, A., Leavitt, B. R., Roos, R. A., Borowsky, B., Landwehrmeyer, B., Frost, C., Johnson, H., Craufurd, D., Reilmann, R., Stout, JC., Langbehn, DR., & TRACK-HD Investigators. (2013). Predictors of phenotypic progression and disease onset in premanifest and early-stage Huntington’s disease in the TRACK-HD study: Analysis of 36-month observational data. The Lancet Neurology, 12(7), 637–649. https://doi.org/10.1016/S1474-4422(13)70088-7

Telezhkin, V., Schnell, C., Yarova, P., Yung, S., Cope, E., Hughes, A., Thompson, BA., Sanders, P., Geater, C., Hancock, JM., Joy, S., Badder, L., Connor-Robson, N., Comella, A., Straccia, M., Bombau, G., Brown, JT., Canals, JM., Randall, AD., Allen, ND., & Kemp, PJ. (2016). Forced cell cycle exit and modulation of GABAA, CREB, and GSK3β signaling promote functional maturation of induced pluripotent stem cell-derived neurons. American Journal of Physiology. Cell Physiology, 310(7), C520–541. https://doi.org/10.1152/ajpcell.00166.2015

Tsvetkov, A. S., Miller, J., Arrasate, M., Wong, J. S., Pleiss, M. A., & Finkbeiner, S. (2010). A smallmolecule scaffold induces autophagy in primary neurons and protects against toxicity in a Huntington disease model. Proceedings of the National Academy of Sciences, 107(39), 16982–16987. https://doi.org/10.1073/pnas.1004498107

Victor, M. B., Richner, M., Olsen, H. E., Lee, S. W., Monteys, A. M., Ma, C., … Yoo, A. S. (2018). Striatal neurons directly converted from Huntington’s disease patient fibroblasts recapitulate age-associated disease phenotypes. Nat Neurosci. doi:10.1038/s41593-018-0075-7

Wang, A. S., Ong, P. F., Chojnowski, A., Clavel, C., & Dreesen, O. (2017). Loss of lamin B1 is a biomarker to quantify cellular senescence in photoaged skin. Scientific Reports, 7. doi:ARTN 15678

Wexler, N. S., Lorimer, J., Porter, J., Gomez, F., Moskowitz, C., Shackell, E., … Project, U. S.-V. C. R. (2004). Venezuelan kindreds reveal that genetic and environmental factors modulate Huntington’s disease age of onset. Proc Natl Acad Sci U S A, 101(10), 3498–3503. doi:10.1073/pnas.0308679101

Wheeler, V. C., Lebel, L. A., Vrbanac, V., Teed, A., te Riele, H., & MacDonald, M. E. (2003). Mismatch repair gene Msh2 modifies the timing of early disease in Hdh(Q111) striatum. Hum Mol Genet, 12(3), 273–281. doi:10.1093/hmg/ddg056

Whittemore, E. R., Loo, D. T., Watt, J. A., & Cotman, C. W. (1995). A detailed analysis of hydrogen peroxide-induced cell death in primary neuronal culture. Neuroscience, 67(4), 921–932. doi:10.1016/0306-4522(95)00108-u

Wu, Y.-Y., Chiu, F.-L., Yeh, C.-S., & Kuo, H.-C. (2019). Opportunities and challenges for the use of induced pluripotent stem cells in modelling neurodegenerative disease. Open Biology, 9(1). https://doi.org/10.1098/rsob.180177

Yang, D., Wang, C.-E., Zhao, B., Li, W., Ouyang, Z., Liu, Z., Yang, H., Fan, P., O’Neill, A., Gu, W., Yi, H., Li, S., Lai, L., & Li, X.J. (2010). Expression of Huntington’s disease protein results in apoptotic neurons in the brains of cloned transgenic pigs. Human Molecular Genetics, 19(20), 3983–3994. https://doi.org/10.1093/hmg/ddq313

Yoo, D. Y., Jung, H. Y., Kim, J. W., Yim, H. S., Kim, D. W., Nam, H., & Hwang, I. K. (2016). Reduction of dynamin 1 in the hippocampus of aged mice is associated with the decline in hippocampal-dependent memory. Molecular Medicine Reports, 14(5), 4755–4760. https://doi.org/10.3892/mmr.2016.5804

Zeron, M. M., Hansson, O., Chen, N., Wellington, CL., Leavitt, BR., Brundin, P., Hayden, MR., & Raymond, LA. (2002). Increased sensitivity to N-Methyl-D-Aspartate receptor-mediated excitotoxicity in a mouse model of Huntington’s disease. Neuron, 33(6), 849–860. https://doi.org/10.1016/S0896-6273(02)00615-3

Zhang, Y., Leavitt, BR., van Raamsdonk, JM., Dragatsis, I., Goldowitz, D., MacDonald, ME., Hayden, MR., & Friedlander, RM. (2006). Huntingtin inhibits caspase-3 activation. The EMBO Journal, 25(24), 5896–5906. https://doi.org/10.1038/sj.emboj.7601445

