## Supplemental figures for "The interaction of aging and oxidative stress contributes to pathogenesis in mouse and human Huntington disease neurons"

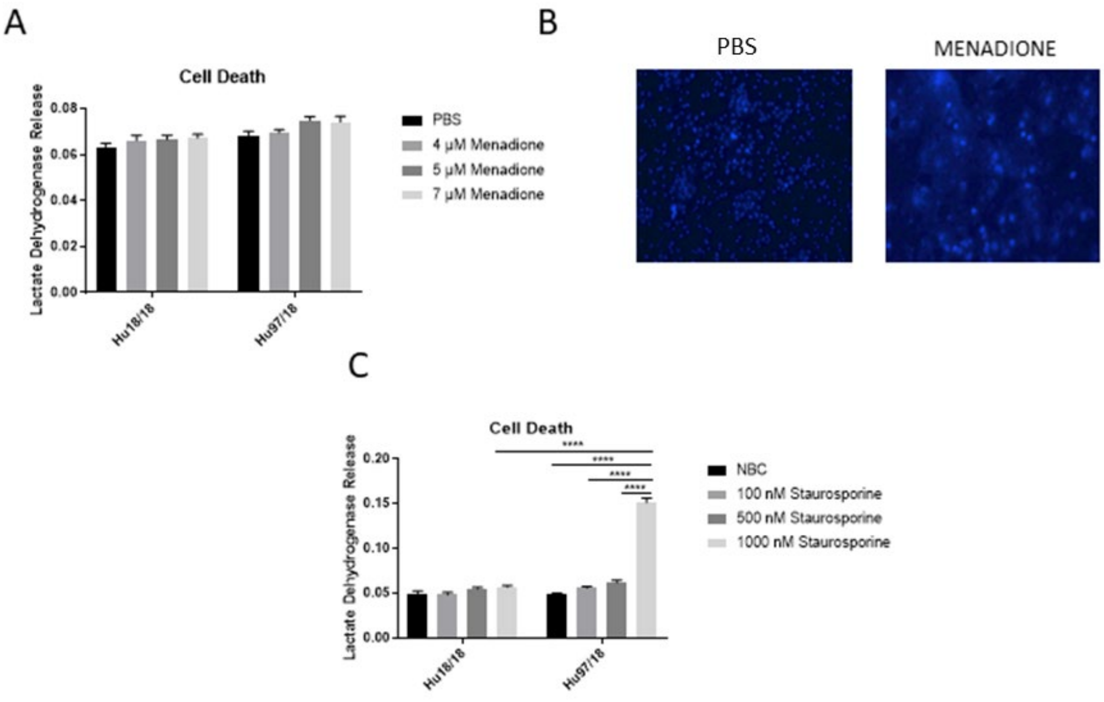


**Figure supplement 1.** (**A**) Menadione treatment does not alter cell death in control or HD neurons. (**B**) Menadione causes DAPI leakage from the nucleus. (**C**) HD neurons are hypersensitive to staurosporine at a high concentration.

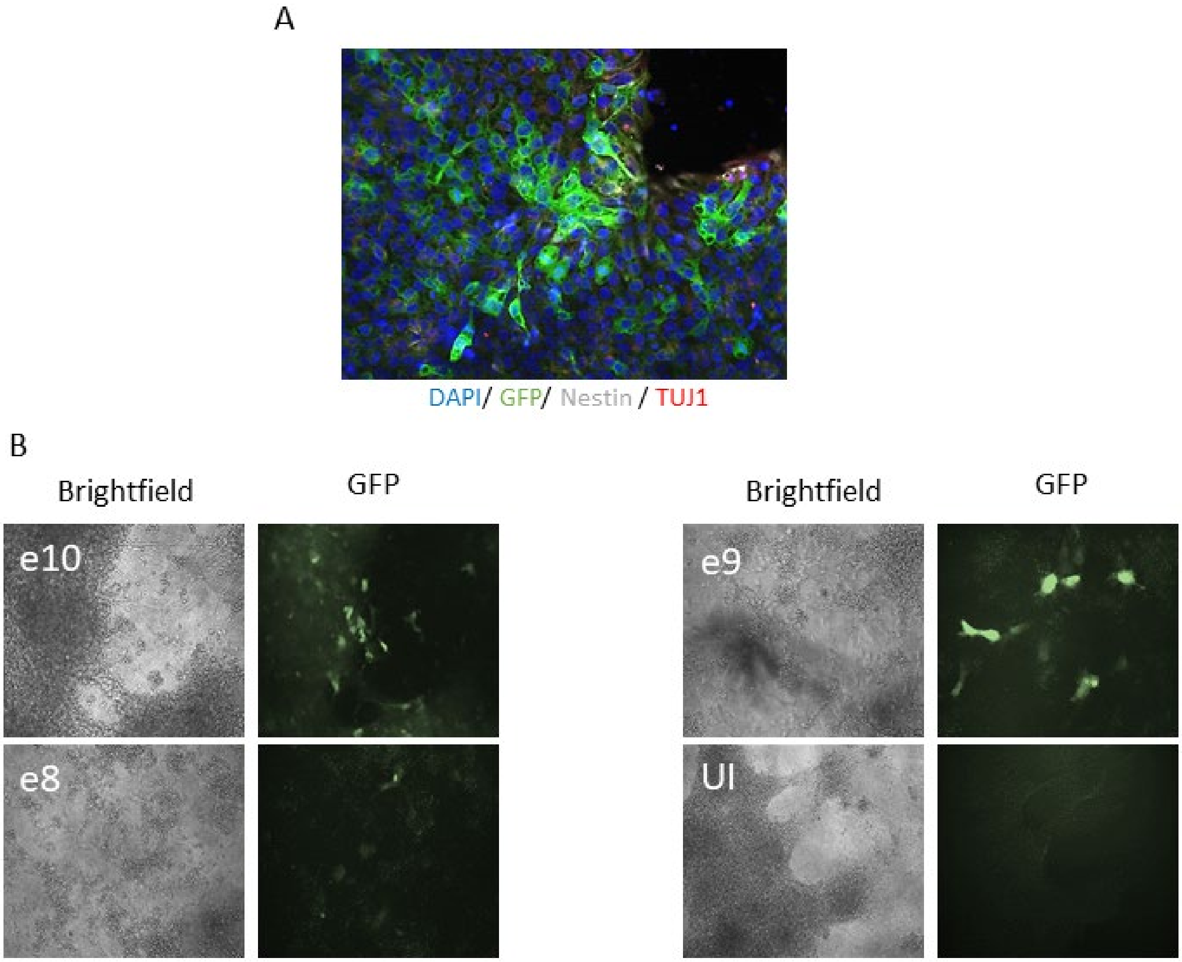


**Figure supplement 2. AAV2/1 GFP infects striatal-like neurons and differentiating iSPC colonies.** (**A**) AAV2/1 GFP infects differentiating iPSC colonies. (**B**) AAV2/1 GFP infects striatal-like neurons differentiated from iPSCs, with highest infectivity at e10.

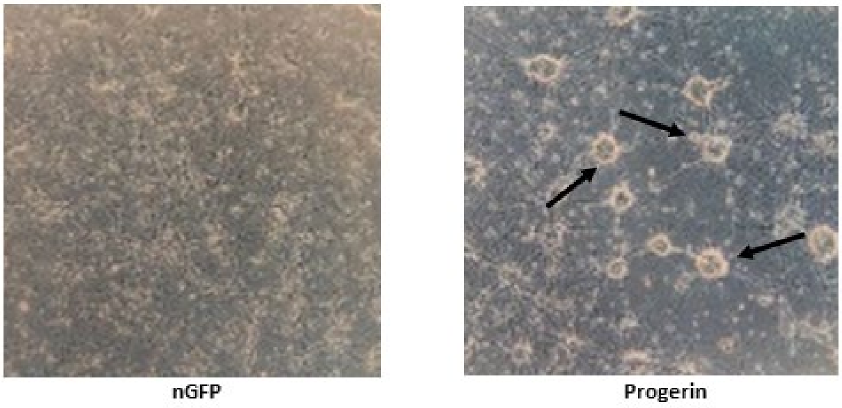


**Figure supplement 3. Cellular rearrangement in HD neurons post-treatment with progerin.**

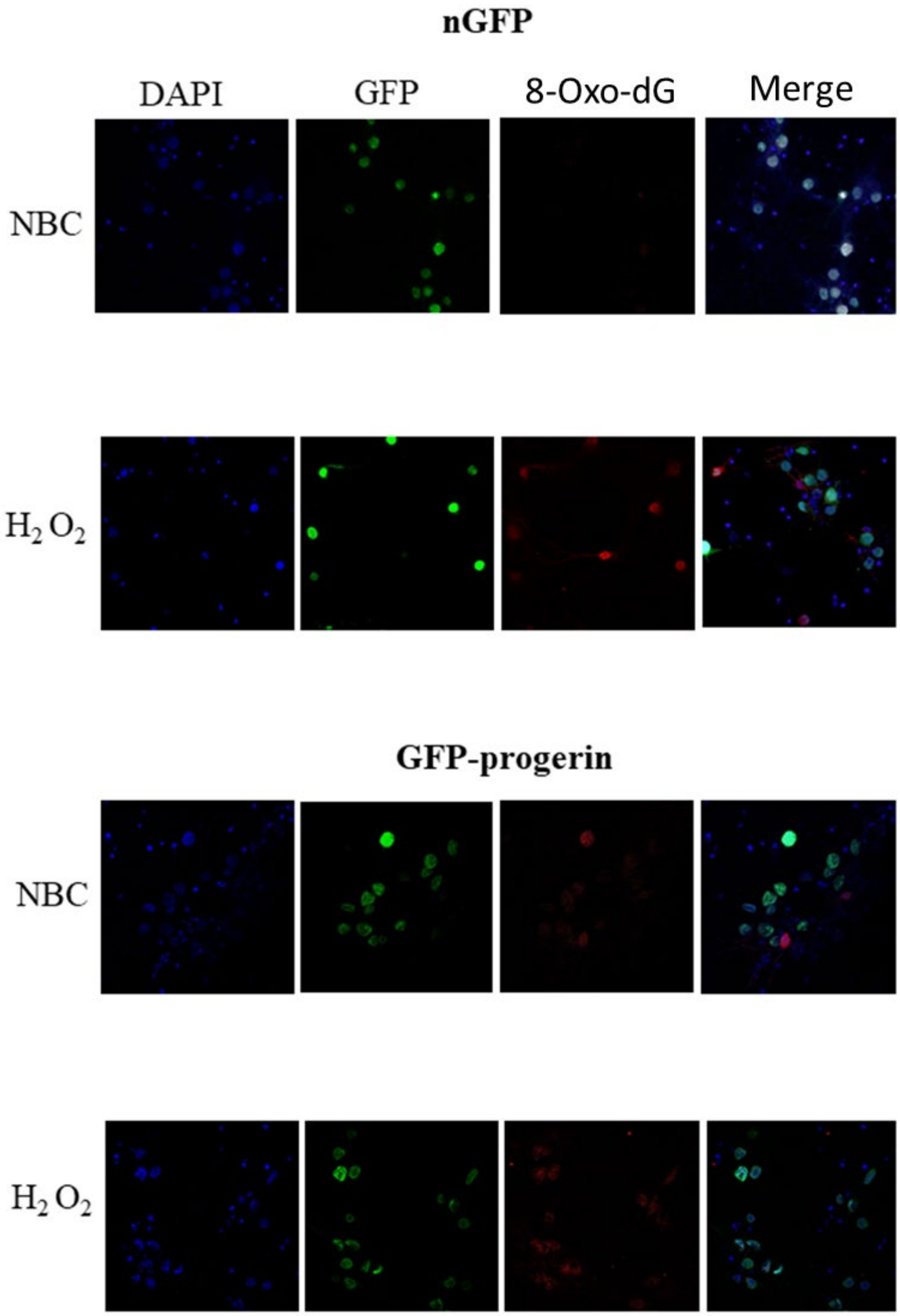


**Figure supplement 4. Representative images of DNA damage in aged Hu18/18 neurons treated with oxidative stress.**

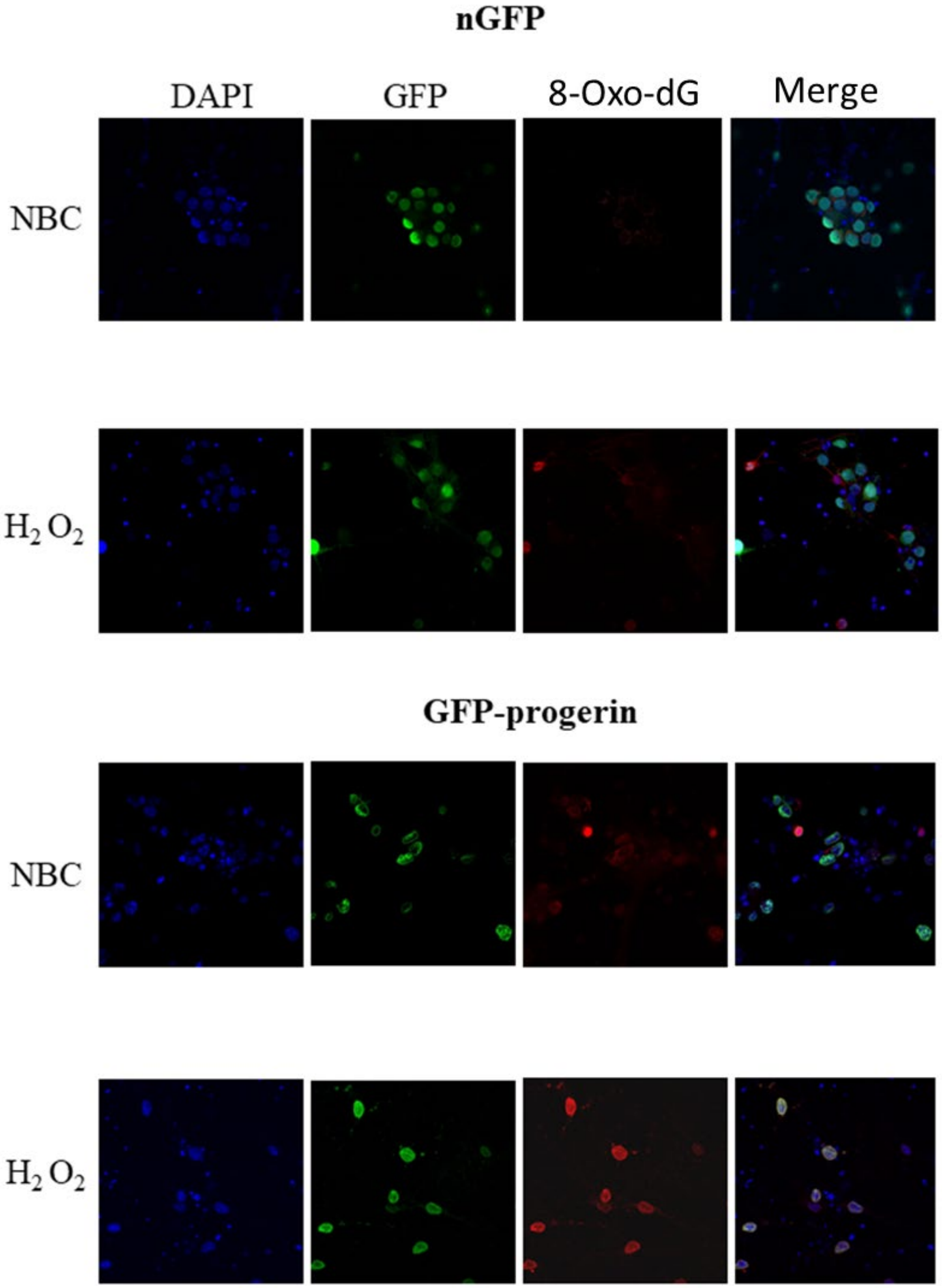


**Figure supplement 5. Representative images of DNA damage in aged Hu97/18 neurons.**

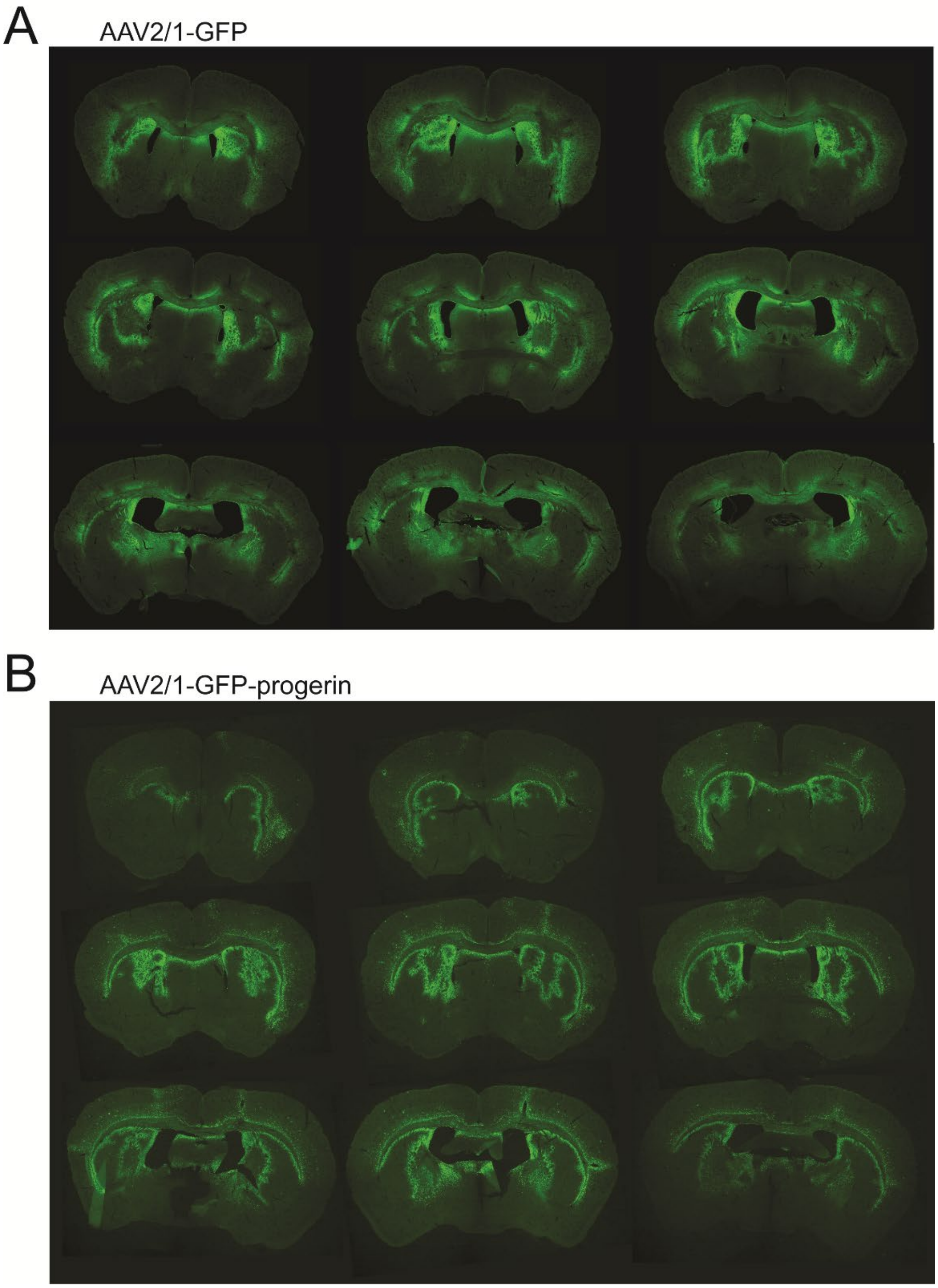


**Figure supplement 6. Distribution of AAV2/1-GFP and AAV2/1-GFP-progerin in the brain of YAC128 mice following bilateral intrastriatal infusion by convection-enhanced delivery at 8 weeks post-injection.**

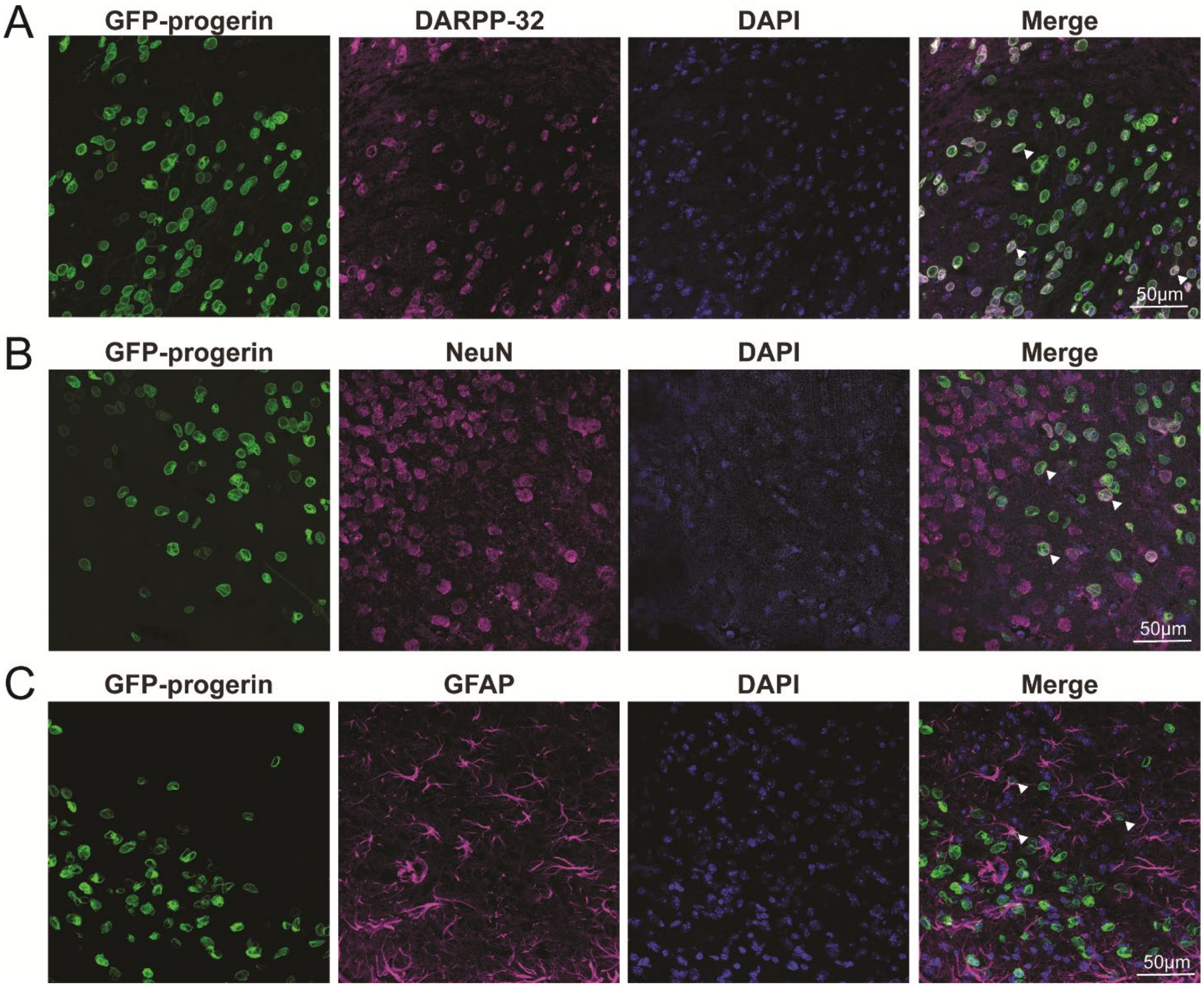


**Figure supplement 7. Tropism of AAV2/1-GFP-progerin in the striatum of YAC128 mice at 8 weeks post-injection.** GFP-progerin transduces (**A**) MSNs, (**B**) neurons, and (**C**) astrocytes/neural progenitors in the striatum. White arrows highlight overlap between GFP-progerin and either DARPP-32, NeuN or GFAP. Scale bar = 50µm.
